# Delineating redox cooperativity in water-soluble and membrane multiheme cytochromes through protein design

**DOI:** 10.1101/2024.03.21.586059

**Authors:** Benjamin J. Hardy, Paulina Dubiel, Ethan L. Bungay, May Rudin, Christopher Williams, Christopher J. Arthur, Matthew J. Guberman-Pfeffer, A. Sofia Oliveira, Paul Curnow, J. L. Ross Anderson

## Abstract

Nature has evolved diverse electron transport proteins and multiprotein assemblies essential to the generation and transduction of biological energy. However, substantially modifying or adapting these proteins for user-defined applications or to gain fundamental mechanistic insight can be hindered by their inherent complexity. *De novo* protein design offers an attractive route to stripping away this confounding complexity, enabling us to probe the fundamental workings of these bioenergetic proteins and systems, while providing robust, modular platforms for constructing completely artificial electron-conducting circuitry. Here, we use a set of *de novo* designed mono-heme and di-heme soluble and membrane proteins to unpick the contributions of electrostatic micro-environments and dielectric properties of the surrounding protein medium on the inter-heme redox cooperativity that we have previously reported. Experimentally, we find that the two heme sites in both the water-soluble and membrane constructs have broadly equivalent redox potentials in isolation, in agreement with Poisson-Boltzmann Continuum Electrostatics calculations. BioDC, a Python program for the estimation of electron transfer energetics and kinetics within multiheme cytochromes, also predicts equivalent heme sites, and reports that burial within the low dielectric environment of the membrane strengthens heme-heme electrostatic coupling. We conclude that redox cooperativity in our diheme cytochromes is largely driven by heme electrostatic coupling and confirm that this effect is greatly strengthened by burial in the membrane. These results demonstrate that while our *de novo* proteins present minimalist, new-to-nature constructs, they enable the dissection and microscopic examination of processes fundamental to the function of vital, yet complex, bioenergetic assemblies.

## 1. Introduction

Multiheme cytochromes are an abundant group of water-soluble and transmembrane proteins that contain multiple redox-active heme cofactors generally held in sufficiently close proximity to facilitate rapid inter-cofactor electron transfer. They are essential to aerobic respiration and play many organism-dependent roles in anaerobic respiration and photosynthesis^1–4^. The biophysical properties and reactivity of these heme cofactors are finely controlled by their conformation^5,6^, axial ligation, local protein environment^7,8^, solvent accessibility and the polarity of the surrounding medium; parameters which have been manipulated and tailored throughout evolution^9^. As the heme cofactors are bound in close proximity, there is an interdependency between their redox properties^10^, resulting from changes in heme-heme electrostatic interactions when an electron is added or removed from one of the pair. This phenomenon, known as redox cooperativity, is thought to promote efficient and rapid directional electron transfer, as observed in the transmembrane cytochrome *bc* ^11^ and *b_6_f* complexes^12^, terminal oxidases^13,14^, cytochrome components of some bacterial photosynthetic reaction centres^15^, water-soluble cytochromes *c*3^16^ and the soluble and transmembrane domains of extracellular heme wires^17^.

Much effort has been made to unpick the specific factors that control the strength of electrostatic coupling and the resulting splitting of heme redox potentials (ΔE_m_) in such proteins^12,18–21^, though the ease of re-engineering these natural cytochromes is severely limited by their inherent complexity. Whilst there have been notable successes in altering the redox potentials of natural membrane cytochromes through heme ligand substitutions (including the *bc*_1_ complex^22,23^ and the decaheme MtrC^24^), most reported mutations confer only minimal effects and often disrupt heme binding sites to the point where the cofactor is lost entirely^25^. In parallel to these experimental efforts, computational studies have been instrumental in elucidating the contributions of electrostatics, electronic coupling and insulation from solvent^1,21,26,27^, providing valuable electrochemical insight into otherwise spectroscopically equivalent hemes that can prove resistant to experimental interrogation^28^.

An alternative, and experimentally tractable, way of studying the fundamental properties of hemes within proteins is through the use of simple ‘maquette’ proteins, which are typically water-soluble or amphipathic four-helix bundles designed to bind redox-active porphyrins^29–32^. Maquettes provide a minimal platform where the protein environment is defined and highly mutable, the sequence is devoid of evolutionary history or complexity, and hemes and related porphyrins can be easily added or removed. When multiple hemes are bound within close proximity, redox cooperativity reminiscent of natural cytochromes is commonly observed^30,33,34^. Whilst the ease of binding porphyrins and other cofactors within maquettes has enabled the design of a multitude of highly functional proteins^35–41^, these have typically lacked a well-defined, singular structure, hindering subsequent engineering with atomistic precision. To address this issue, we designed a suite of structured, robust and modular coiled-coiled *de novo* heme proteins which are readily expressed in *E. coli*^42^.

This family of redox proteins is based upon the diheme protein called 4D2, which binds two bis-histidine coordinated *b*-type hemes, for which a crystal structure was solved to 1.9 Å resolution^33,42^. The two co-planar hemes sit in near-identical binding sites and exhibit noticeable redox cooperativity, resulting in a ΔE_m_ of 63 mV. From 4D2 we generated a rigid monoheme protein, m4D2, by removing one heme binding site and repacking the vacant heme binding site through computational protein design. 2D NMR spectroscopy revealed m4D2 to be well-structured when loaded either with heme *b* or a symmetrical heme analogue. By duplicating the sequence of each helix of 4D2 we also generated an extended tetraheme cytochrome, e4D2, which could be directly observed by electron microscopy despite its small size of 24 kDa. To study the properties of the diheme scaffold within the membrane, we used hydrophobic surface design^43^ to create a transmembrane version of 4D2, termed CytbX, which was efficiently routed to bacterial membranes where it innately bound two *b*-type hemes with high affinity. Interestingly, when bound within the transmembrane CytbX, the heme pair exhibits significantly stronger redox cooperativity than 4D2, manifesting a larger ΔE_m_ of 113 mV despite the interior heme binding sites being nearly identical to those in 4D2.

With the successful 4D2 and CytbX designs in hand, we are in the unique position to systematically study the interactions between a pair of *b*-type hemes within the same soluble protein scaffold, and directly compare them with the membrane equivalent. While it has not been possible to obtain a structure for CytbX, all experimental observations are consistent with 4D2 and CytbX adopting the same overall fold, inter-heme distance, relative heme orientations and internal heme-binding residues. These similarities rule out many confounding factors that can modulate the ΔE_m_ of the heme pair. Our approach, therefore, provides a valuable reductionist system in which to directly study the effect of the membrane on heme redox potentials and redox cooperativity.

To this end, we report here the design, cellular production, and characterisation of a full set of single-heme 4D2 and CytbX constructs. We exploit the modular nature of our coiled-coil system to swap the heme-binding and core packing modules of m4D2, yielding the complementary soluble monoheme construct. We also apply the same computational design pipeline to CytbX to yield two transmembrane monoheme cytochromes. Experimentally, we observe that the local environments of each heme binding site contribute minimally to the ΔE_m_ observed in the diheme constructs and, using two independent computational methods for redox potential prediction, we conclude that redox potential splitting in our system is mainly driven by heme-heme electrostatic interactions. Furthermore, redox potential calculations support the observation that the strengthened redox cooperativity in CytbX is indeed a function of the surrounding low dielectric medium of the membrane (or detergent micelle) and enable us to assign the low and high potential hemes of 4D2 and CytbX. Our findings demonstrate that both *de novo* soluble and membrane cytochromes can be engineered with atomic precision and demonstrate the utility of our platform for studying the fundamental properties and interactions of hemes within proteins.

## 2. Results

### 2.1 Protein Design

We previously reported the design of the monoheme protein m4D2 (Figure 1a) from the diheme 4D2 (Figure 1b) by retaining one of the heme binding sites (heme 1) whilst repacking the other site (heme 2) to create a compact hydrophobic core^42^. By retaining heme 1, we sought to preserve hydrogen bonding interactions observed in the 4D2 and 4D2-T19D crystal structures (PDB IDs: 7AH0, 8CCR) between the heme propionate groups and sidechains of the nearby loops. Here, we wished to design the complementary or inverse protein, with heme 2 preserved and heme site 1 re-packed. To distinguish between these constructs, we term the monoheme designs retaining heme in sites 1 and 2 as m1-4D2 (previously m4D2) and m2-4D2 respectively (Figure 1c). To design the sequence of m2-4D2, we simply swapped the order of the heme-binding and hydrophobic packing portions of each helix of m1-4D2 (Figure 1a,c and Figure S17) without any additional modifications.

**Figure 1.**
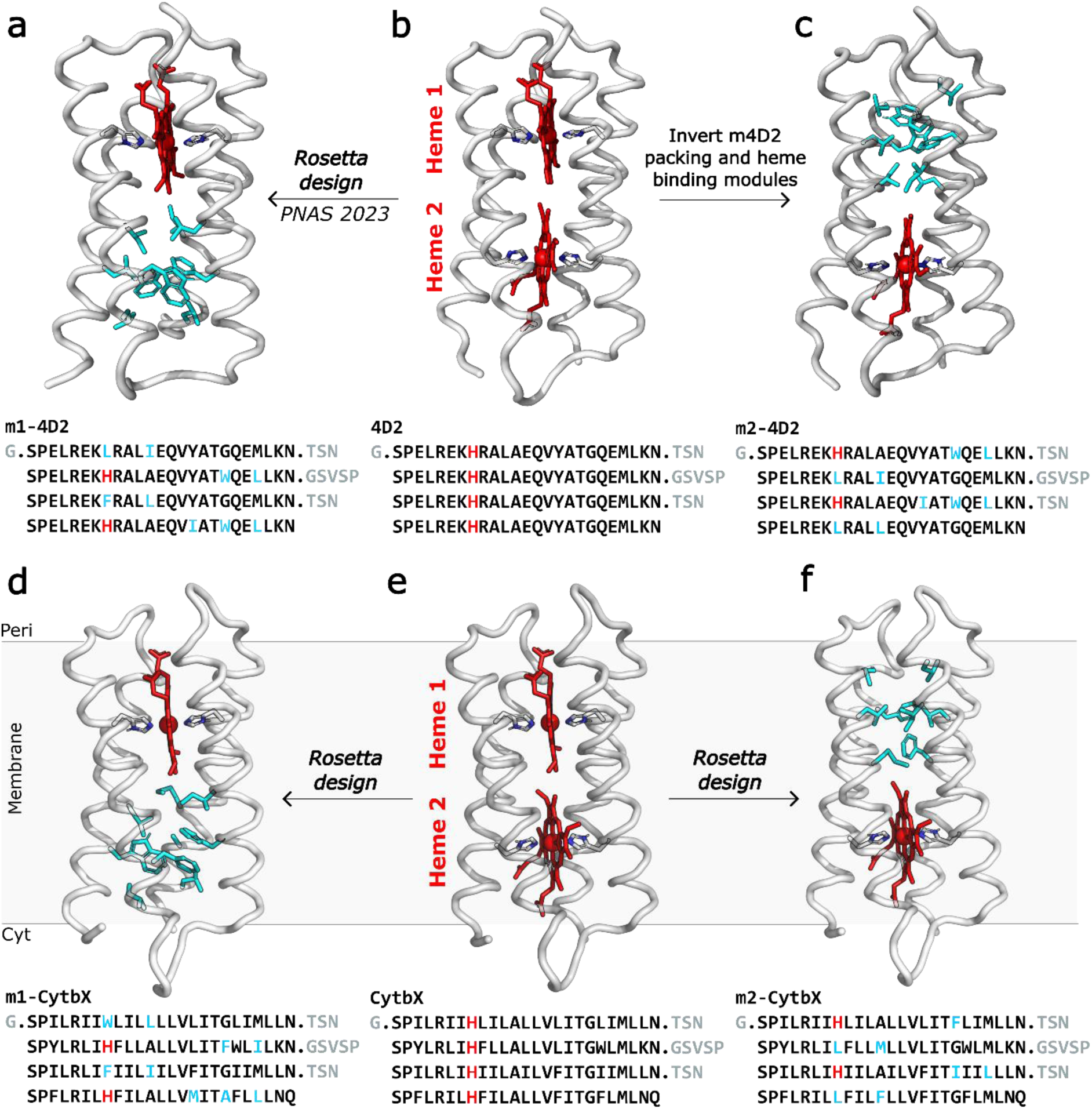
The suite of monoheme and diheme water-soluble and transmembrane modular redox proteins based on the coiled-coil diheme protein 4D2. (**a**) m1-4D2 contains heme 1 but heme site 2 is re-designed. (**b**) 4D2 is the basis of the protein suite and contains two identical bis-histidine heme binding sites. (**c**) m2-m4D2 contains heme 2 but heme site 1 is re-designed, containing the same core packing residues as in site 2 if m1-4D2. (**d**) m1-CytbX contains heme 1 of CytbX but heme site 2 was re-designed with Rosetta. (**e**) CytbX is the transmembrane version of 4D2. (**f**) m2-CytbX contains heme in site 2 but heme site 1 was re-designed with Rosetta. All structures are Rosetta-generated models, apart from 4D2 which is the crystal structure (PDB ID: 7AH0) with reconstructed loops. Protein backbones are shown as grey ribbons, hemes (red), mutated core residues (cyan) and heme-coordinating residues (grey) are shown as grey sticks. The sequence of each protein is shown, with mutated residues highlighted in cyan, histidines highlighted in red, and loop residues in grey.

We generated a theoretical model of m2-4D2 by threading the new sequence onto the crystal structure of 4D2 (containing a reconstructed loop) using Rosetta^44^, and subjected this model to structural relaxation (Cα RMSD vs 4D2 = 1 Å). In this model, the swapped modules form a heme-binding pocket and compact core as desired (Figure 1c). ESMfold^45^ predicts a high confidence structure (mean pLDDT = 87.0) that mostly matches the theoretical model (Cα RMSD = 2.63 Å, TM = 0.73; see Table S2, Figure S5), containing a heme-shaped cavity in heme site 2, with histidine side chains poised for heme coordination, and a compact core in the packing module (Figure S1a, b). AlphaFold2 (AF2) predicts a high confidence structure (mean pLDDT = 82.3) containing a widened heme-shaped cavity in heme site 2 and compact core in heme site 1 (Figure S1c, d), though it predicts the helices of m2-4D2 to pack in an incorrect, mirrored topology (Figure S2).

Interestingly, AF2 also predicts this same mirrored helical topology for many of our *de novo* heme proteins, including m1-4D2 and CytbX, and most notably for 4D2, where two crystal structures have been solved in the expected topology (Figure S3d). For related proteins such as m1-4D2, AF2 predicts no heme-shaped cavity at all (Figure S4). In stark contrast, the ESMfold structure prediction of 4D2 aligns almost exactly to the crystal structure (Cα RMSD = 1.24 Å, TM = 0.92), with correct helical packing topology, heme-shaped cavities, and with heme-coordinating histidine-threonine H-bonding pairs superimposable (Figure S3a-c). We, therefore, suggest a general suitability of ESMfold over AF2 for the structure prediction of *de novo* heme proteins, in agreement with other published observations that ESMfold outperforms AF2 on amino acid sequences with no homology to natural proteins^46^. ESMfold structure predictions of the complete suite of proteins discussed in this report are shown in Figure S5, and prediction metrics from ESMfold and AF2 are compared in tables S1 and S2.

To generate single-heme CytbX variants, we used RosettaMP^47^ with the *franklin2019*^48^ membrane protein score function to computationally sample mutations for repacking either heme site. Here, we refer to CytbX with heme 1 present as m1-CytbX (Figure 1d) and heme 2 present as m2-CytbX (Figure 1f), these being the transmembrane counterparts of m1-4D2 and m2-4D2 respectively (Figure 1a, c). We tested whether the m1-4D2 core residues could be directly transplanted into CytbX but found that these would likely splay the bundle (Figure S6) (see Methods). Using a flexible-backbone design protocol, we allowed substitutions (to any hydrophobic amino acid) at these residue positions in CytbX to obtain optimised hydrophobic cores. Given that the positions were originally selected to not alter any knobs-into-holes (KIH) packing residues, they do not disrupt vital transmembrane helix-helix interactions in CytbX^42,43^. Decoys were ranked by their Rosetta score and the top 10 sequences for each design were further analysed. Multiple sequence alignment with ClustalOmega^49^ revealed redundancy in the top sequences (Figure S7). For both constructs, we selected unique sequences that had either the lowest Rosetta score or the most compact core (as reported by the Packstat^50^ metric) for bacterial expression (Figure S7). While all four designs (two designs for each of two constructs) were expressed and purified from *E. coli* membranes, exhibiting identical biophysical properties, only the best expressing sequences from each pair (Figure S11a) for m1-CytbX and m2-CytbX are discussed further for clarity.

The Rosetta-generated models of m1-CytbX and m2-CytbX contained compact hydrophobic cores (Figure 1d, f) and minimal structural deviations from the CytbX backbone (M1-CytbX Cα RMSD = 1.02 Å, m2-CytbX Cα RMSD = 0.96 Å). Both designs were predicted to contain four TM helices with N_in_-C_in_ topology by the DeepTMHMM server (Figure S10)^51^. ESMfold predicted very high confidence structures closely matching the Rosetta designs (Figure S5) containing heme-shaped cavities and compact hydrophobic cores at the opposite ends of the bundles (Figure S8). Again, AF2 predicted lower-confidence, incorrectly packed structures lacking heme-binding cavities (Figure S9).

### 2.2 Expression and characterisation of mono-heme proteins

We subsequently ordered codon-optimised synthetic genes encoding the designed amino acid sequences for m2-4D2, m1-CytbX and m2-CytbX in standard pET expression vectors (pET-151, pET-29 or pET-21) with TEV (Tobacco etch virus N1A) protease-cleavable *N*-terminal hexahistidine or thrombin-cleavable Strep tags for the soluble and membrane proteins respectively. Following expression in *E. coli* T7-express cells, we purified the soluble m2-4D2 using affinity chromatography procedures as previously described^42^, cleaved the purification tag with TEV protease, loaded the incompletely heme-incorporated m2-4D2 with exogenous heme, and purified the holoprotein by size-exclusion chromatography (Figure S11). In contrast, we expressed the m1-CytbX and m2-CytbX membrane proteins in *E. coli* C43 (DE3) and used Strep-Tactin affinity chromatography to purify the proteins from isolated membranes, solubilised using the mild non-ionic detergent CYMAL-5. Following tag removal with Thrombin protease, we used SEC with no further addition of heme. Both m1-CytbX and m2-CytbX eluted as single monodisperse species, whilst m2-4D2 eluted as two sharp peaks (Figure S11)

All purified proteins exhibited an intense red colour with UV-visible absorbance spectra containing prominent ferric Soret peaks at 416-417 nm consistent with bis-histidine coordination of heme *b* ^42,43,52^ (Figure 2 a-c). Unexpectedly, the absorbance spectrum of m2-CytbX contained an additional low intensity peak in the Q-band region at 590 nm characteristic of zinc protoporphyrin IX (ZnPPIX)^53^ (Figure 2c), and a broadened Soret peak at 417 nm. The presence of ZnPPIX was confirmed by fluorescence spectroscopy and was estimated to occupy fewer than 10% of all binding sites (Figure S12). The binding stoichiometry of all three monoheme designs was confirmed as 1:1 protein:heme by native mass spectrometry (MS) (Figure S13, Table S3). The small portion of ZnPPIX-bound m2-CytbX could not be identified in the MS spectrum.

**Figure 2.**
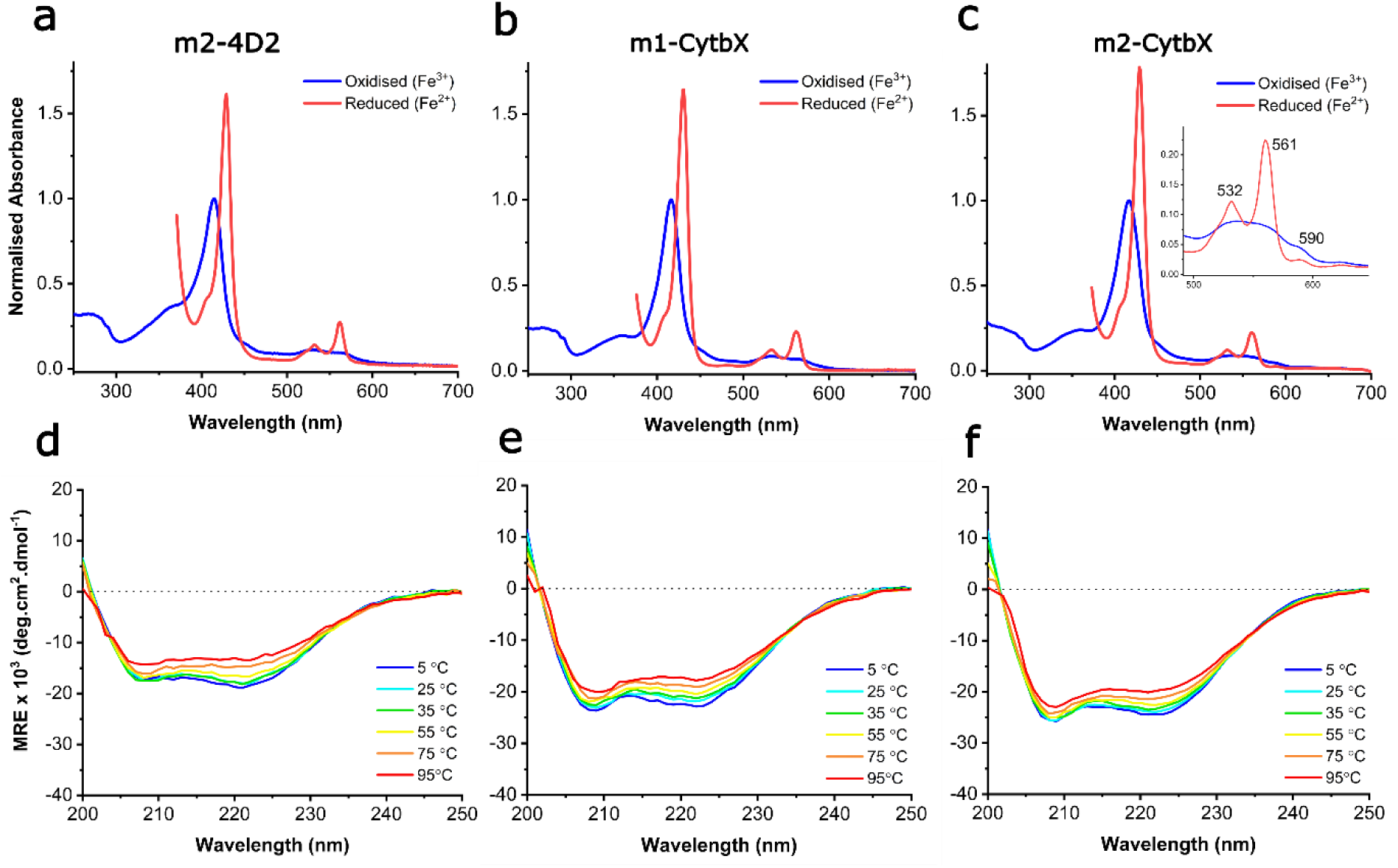
Biophysical characterisation of monoheme designs. UV-Vis absorbance spectra of purified (**a**) m2-4D2, (**b**) m1-CytbX and (**c**) m2-CytbX. Reduced spectra are cropped at the point where the detector is saturated by dithionite absorbance. Absorbance is normalised to the oxidised Soret peak maximum. Circular dichroism spectra during thermal melts of purified (**d**) m2-4D2, (**e**) m1-CytbX and (**f**) m2-CytbX from 5 °C to 95 °C. All designs bind heme, are helical and highly thermostable as designed.

We then used far-UV circular dichroism (CD) spectroscopy to confirm that the new holo-proteins were predominantly helical, with all exhibiting remarkable thermostability (T_m_ > 95 °C) (Figure 2 d-f). The CD spectra exhibited a small, linear decrease in MRE at 222 nm up to 95 °C with no observable melt transition (Figure S14). The CD spectrum of m2-4D2 reveals a higher ε_222nm_/ε_208nm_ than both m1-CytbX and m2-CytbX (Figure 2d), suggesting a more supercoiled structure^54^. Whilst crystal structures for these three proteins could not be obtained, the H^1^-N^15^ TROSY spectrum of m2-4D2 shows similarly good peak dispersion to that of m1-4D2, indicating that it is well structured in the heme-bound state (Figure S15). This confirms that combining the heme-binding and packing modules of m1-4D2 in either orientation and in such a simple, modular fashion, yields well-structured holo-proteins.

### 2.3 Redox properties of water-soluble and transmembrane monoheme and diheme proteins

To measure the redox potentials (E_m_) of the *b*-type hemes bound within these proteins, we employed optically transparent thin layer electrochemistry (OTTLE)^55^. Potentiometric titrations revealed a single E_m_ for m2-4D2 at -125 ± 2 mV (vs NHE) (Figure 3a), very close to the previously-measured E_m_ for m1-4D2 of -117 mV^42^, suggesting that the hemes of m1-4D2 and m2-4D2 reside within similar chemical environments despite residing at opposing ends of the helical bundle. These potentials are closer to the high-potential heme of 4D2 (-104 mV) than the low-potential heme (-167 mV), and the difference between these potentials (8 mV) represents approximately 13% of the redox splitting measured for the 4D2 hemes (63 mV) (Table S4). This strongly suggests that the redox splitting in 4D2 is mainly a result of heme-heme interactions rather than differences in heme environments.

**Figure 3.**
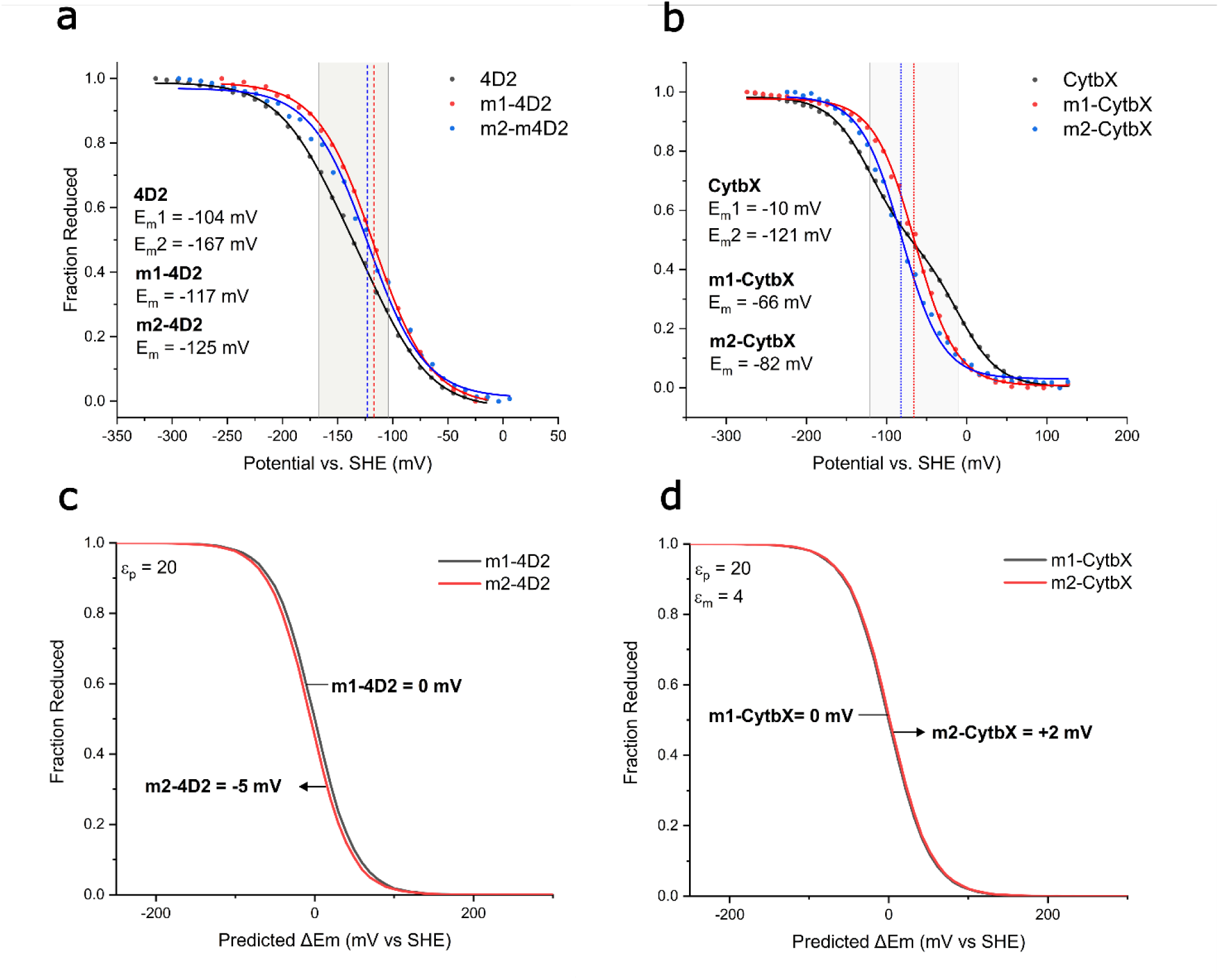
Redox potentiometry of the monoheme and diheme (a) membrane proteins and (b) water soluble proteins. Measurements were performed with purification tags removed from all proteins. Membrane proteins: pH = 7.4, soluble proteins: pH = 8.6. Buffer for membrane proteins: 50 mM NaP (pH 7.4), 150 mM NaCl, 5% glycerol, 0.08% CYMAL-5. Poisson-Boltzmann Monte-Carlo electrostatics calculations of redox titrations for (c) m1-4D2 vs. m2-4D2 and (d) m1-CytbX vs. m2-CytbX. Water-soluble proteins were modelled in a solvent with a dielectric constant (ε_p_) of 20 at pH 8.6, and membrane proteins were modelled in an implicit membrane with ε_m_=4 at pH 7.4. Predicted E_m_ shifts are reported using m1-4D2 and m1-CytbX as references for m2-4D2 and m2-CytbX respectively. Redox potentials were derived by fitting to a Nernst equation describing a single electron transfer.

Potentiometric titrations revealed single E_m_s for m1-CytbX and m2-CytbX of -66 ± 2 mV and -82 ± 1 mV respectively, with both residing between the two split potentials of diheme CytbX (-10 mV and -121 mV) (Figure 3b). The small portion of ZnPPIX bound to m2-CytbX does not contribute to the potentiometric data as ZnPPIX is not redox active within the window of potential scanned in these measurements. The difference in redox potentials of the single-heme membrane proteins (16 mV) is double that of the soluble counterparts (8 mV) and represents about 5% of the total redox difference of CytbX (111 mV). As for 4D2, these findings suggest that heme-heme interactions, rather than differences in heme environment, dominate the overall ΔE_m_ in CytbX. The increased magnitude of ΔE_m_ in CytbX versus 4D2 suggests a predominantly electrostatic origin for the split in heme *b* redox potentials, as the lower dielectric environment of the membrane protein interior would less efficiently screen the electrostatic field felt by each heme.

To further probe the electrostatic contribution to the redox cooperativity, we calculated electrostatic maps and isosurfaces for each of the soluble and membrane proteins. These revealed that m1-4D2 and m2-4D2 have non-identical distributions of charge densities (Figure S16b and c), likely accounting for the small differences in their redox potentials. Of particular note is a smaller density of negative charge near the heme of m2-4D2 (Figure S16 c), and a greater density of positive charge near the heme of m1-4D2 (Figure S16 b), relative to 4D2. This highlights that the recombination of individual modules in these new, swapped constructs alters electrostatic properties, and that such changes should be assessed during design. The computed electrostatic isosurfaces also highlight the strong enrichment of positive charge on the cytoplasmic side of CytbX, a direct result of implementing the positive-inside rule to enforce N_in_-C_in_ transmembrane topology during design (Figure S16 d-f)^43,56,57^, likely influencing the potential of heme 2.

Given the experimental results, we wished to further investigate the influence of electrostatics on the heme potentials of the full suite of 4D2 proteins using two computational methods: (i) an established Poisson-Boltzmann Monte-Carlo (PB-MC) continuum-electrostatic workflow^58–60^ and (ii) BioDC, a new Python program developed to model redox cooperativity and conductivity in multi-heme cytochromes.^1^

As inputs to both computational methods, we used the crystal structure of 4D2 (PDB ID: 7AH0) with reconstructed loops and Rosetta-relaxed models of the five other proteins. The PB-MC calculations predicted a relative shift in E_m_ of -5 mV for m2-4D2 relative to m1-4D2 (Figure 3c), in excellent agreement with the measured difference of -6 mV (Figure 3a). For the membrane proteins, the PB-MC calculations predicted a relative shift of +2 mV for m2-CytbX vs. m1-CytbX (Figure 3d), whilst we experimentally measured a difference of -16 mV (Figure 3b). The positive shift predicted by PB-MC (although minimal) is consistent with the expected effect of enriched positive charge near heme 2 in m2-CytbX. Whilst the reason for the discrepancy between the predicted and measured values is not fully understood, it is likely to be related to the absence of protein dynamics in our PB-MC calculations. Adding conformational protein dynamics (e.g. using molecular dynamics-derived snapshots) to the current PB-MC method should improve the agreement with experimental values. This discrepancy may also suggest that m2-CytbX is adopting a subtly different structure than predicted. Overall, PB-MC predicts the heme site 1 and 2 environments in the water-soluble and membrane proteins to be mostly equivalent, in agreement with experimental measurements.

BioDC predicts the heme redox potential of m2-4D2 to be essentially the same as for m1-4D2, but the potential of m2-CytbX to be shifted by -37 mV relative to m1-CytbX. Both predictions are in good agreement with the experimental observations of -5 and -16 mV shifts (Figure 4).

**Figure 4.**
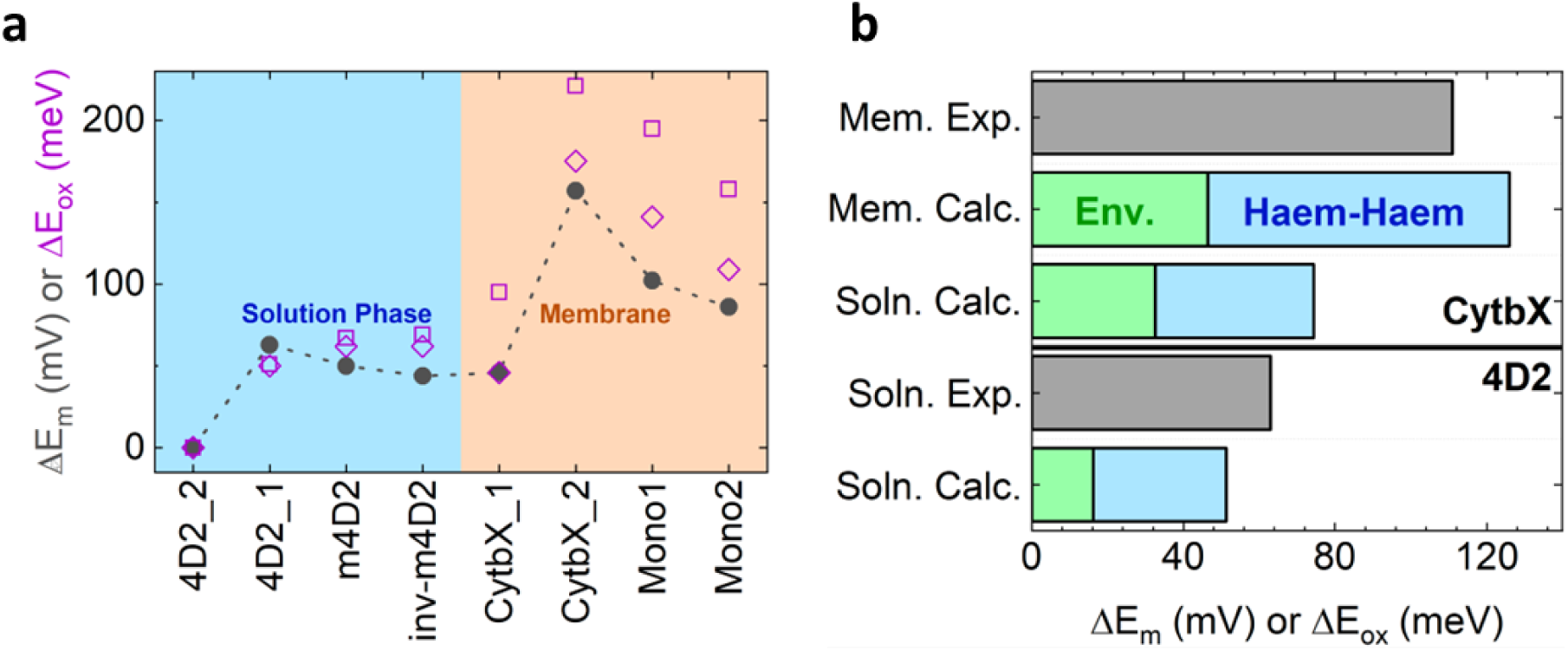
(*a*) BioDC reproduces the trend in redox potential shifts across the mono- and diheme constructs in aqueous and membranous environments, and (*b*) delineates the contributions to the redox potential splitting in the dihemes in terms of environmental electrostatics and heme-heme interactions. Hemes in site 1 and 2 of the diheme constructs are denoted by suffixes (e.g. 4D2_1 denotes 4D2 heme 1). Note that the relative redox potentials for the dihemes from experiment were assigned to heme site 1 and 2 based on the BioDC results. The purple diamonds and squares correspond to results obtained using atomic partial charges for the heme derived according to the Restrained Electro-Static Potential (RESP) and Charge Method 5 (CM5) schemes, respectively.

In the parent diheme constructs, BioDC predicts that heme 2 of 4D2 in an aqueous environment (dielectric constant = 78.2) is shifted by -51 mV relative to heme 1, in good agreement with the 63 mV ΔE_m_ measured experimentally. Heme 1 of CytbX in a membranous environment (dielectric constant = 6.4) is predicted to be shifted by -126 mV relative to heme 2, in good agreement with the 111 mV ΔE_m_ measured experimentally. Interestingly, heme 2 is predicted to have a more positive E_m_ than heme 1 in the diheme CytbX, whereas the heme in m2-CytbX is predicted to have a more negative E_m_ than the heme in m1-CytbX.

The splitting of redox potentials in the diheme 4D2 and CytbX proteins reflects two contributions: (1) The different electrostatic interactions exerted by the environment on each heme while each is in the reduced state, and (2) the electrostatic destabilisation of the oxidised state of one heme by the oxidation of the adjacent heme. Approximately 16 mV (32%) and 46 mV (37%) of the ΔE_m_ of 4D2 and CytbX, respectively, originate from the environmental effect. The change in heme-heme interaction energy upon oxidation of one of the hemes contributes the remaining 35 mV (69%) and 80 mV (63%) mV to the ΔE_m_. All of these results from BioDC are insensitive to the choice of two different schemes for assigning atomic partial charges (RESP and CM5) to the heme in the reduced and oxidised states (Figure 4).

The enlarged potential splitting (ΔE_m_) in CytbX versus 4D2 is attributable to the lower dielectric screening from the environment. Additional BioDC calculations revealed that if CytbX was hypothetically in the same aqueous environment as 4D2, the environment-induced and adjacent-heme-oxidation contributions to the ΔE_m_ decrease from 46 to 33 mV, and 80 to 42 mV, respectively (Table S5), resulting in a ΔE_m_ of 75 mV. This demonstrates that the ΔE_m_ amplification is a result not just of the hydrophobic surface of CytbX but of the embedding of CytbX within a hydrophobic environment.

## 3. Discussion

This study highlights the utility of our robust and highly engineerable redox protein platform for probing redox cooperativity in multiheme cytochromes. We demonstrate that single-heme cytochromes can be readily designed from our water-soluble and transmembrane diheme constructs, and that these retain the rigidity, high affinity heme-binding, and biocompatibility of their parent proteins. Through a combination of experimental and computational approaches, we find that electrostatic coupling between hemes in the diheme 4D2 and CytbX proteins is principally responsible for the observed split in redox potentials, with the differences in the electrostatic micro-environments of the binding sites accounting for the remaining separation in their redox potentials. Importantly, we find that the larger ΔE_m_ observed in the membrane-soluble CytbX is a function of the low dielectric environment present in micelles, and, by extension, the membrane. This environment serves to strengthen the heme-heme electrostatic interactions leading to more pronounced redox cooperativity.

It is likely that this effect is similarly responsible for the ubiquitous split of redox potentials, typically on the order of 100 mV or more, seen in natural multiheme transmembrane cytochromes^22,61^. Functionally, this ΔE_m_ provides an energetically favourable driving force for directional electron transfer from the low (*b*_L_) to high potential (*b*_H_) hemes across the membrane in some, but not all, respiratory complexes^22^. In fact, enzymatic function can be retained in cytochrome *bc*_1_ mutants with no overall difference in heme potentials, while other complexes with similar transmembrane heme proteins can tolerate completely endergonic heme configurations while maintaining function^22^. Whatever the functional implications of the redox split are to overall enzymatic function, our findings here suggest that the magnitude of the diheme redox potential split is likely a direct result of proximal heme pairs being embedded within the hydrophobic environment of the membrane. In addition to acting generally as an electrically insulating medium, the membrane likely plays a role in amplifying electrostatic effects in transmembrane bioenergetic proteins, while also manipulating the redox potentials of embedded redox cofactors through membrane potential^22,61^. Furthermore, it is also known that the dielectric properties of the protein itself can exert fine control over heme properties within the membrane environment. For example, the balance between ΔE_m_ and the dielectric heterogeneity within the dimer interfaces of *bc*_1_ and *b*_6_*f* complexes modulates the extent and efficiency of cytochrome *b* intra-monomer (or cross-branch) electron transfer^19,20^.

In our recent report describing the design of 4D2^42^, we commented that, whilst we observed split heme potentials, we could not assign which were the high and low potential hemes. Now, based upon the BioDC calculations, we suggest that heme 2 and heme 1 are the low and high potential hemes (*b*_L_ and *b*_H_) of 4D2 respectively, whereas heme 1 and heme 2 are *b*_L_ and *b*_H_ in CytbX. Assignment of the low and high potential hemes of CytbX has implications for the favoured direction of electron transport across the membrane. When expressed in the cell, the N_in_-C_in_ topology of CytbX (and monoheme variants) enforced during design dictates that electron transport would be most favourable from the periplasm to the cytoplasm. When assembling CytbX into proteoliposome systems and artificial electron transport pathways more generally, the orientation of the protein in the membrane should therefore be considered for most efficient transmembrane electron transport, in addition to the overall driving force from the redox pair of the terminal electron donors and acceptors.

The construction of the CytbX-Mono proteins here now demonstrates that transmembrane cytochromes that bind only a single heme *b* are accessible by design and provides two such proteins as blank slates for further construction and engineering. It is interesting to observe that the remarkable thermostability of CytbX remains in the CytbX-Mono proteins, despite losing the contributions of two histidine-heme ligations in the removed heme site. This strongly suggests that the Rosetta-designed cores of these proteins are well packed and highly stable. Here we have demonstrated that the modularity of our system enables the simple and facile swapping of heme-binding and packing modules to interrogate heme properties, a feat which is not achievable in natural bioenergetic complexes, and that our diheme protein scaffold is highly engineerable in both its water-soluble and membrane-soluble forms.

## 4. Materials and Methods

### 4.1 Membrane protein design

A Rosetta-generated relaxed model of CytbX was used as a starting point for the design of the CytbX-Mono proteins. To generate mono-heme models in each topology, either of the two hemes were deleted from the CytbX model in PyMol. Residues facing this cavity were then mutated to produce a tightly packed protein core. These residues were taken as the same mutable residues specified in the design of m1-4D2 from 4D2. To generate transmembrane single-heme constructs, we first tested whether core residues of the m1-4D2 packing module could be directly transplanted into CytbX, given that it adopts the same overall backbone structure as 4D2. As an initial test, we copied the m1-4D2 core residues into the corresponding heme site in CytbX (site 2). These mutations were H9L, A13I, G48W, M51L, H68F, A71L, F103I, G106W and M109L.

ESMfold predicts a high-confidence structure (mean pLDDT = 86.3) with minimal deviation to CytbX (Cα RMSD = 2.22 Å) and a heme-shaped cavity in the binding site. However, the m1-4D2 packing residues structurally perturb the bundle, slightly levering the helices apart (Figure S6a, b). When docked with heme and relaxed in Rosetta, the structure adopts a more compact conformation (1.64 Å Cα RMSD vs. CytbX, 2.05 Å Cα RMSD vs. m1-4D2). When the nine m1-4D2 mutations are introduced into the CytbX model through flexible-backbone Rosetta design, generated decoys have 0.7 Å mean Cα RMSD vs. CytbX and compact cores. Although this sequence could adopt suitable structures after Rosetta relaxation, the ESMfold prediction suggested that the sequence could be optimised further.

To obtain optimised repacked cores for both monoheme CytbX structures, we turned to Rosetta design. Mutable residues as outlined above were allowed to mutate to any hydrophobic amino acid (FAMILYVW), defined in a resfile. Models of CytbX with either heme deleted were used as starting structures, and histidine coordination of the remaining heme was enforced using a constraints file during all steps. Design was performed using a custom RosettaScripts XML file, implementing a FastDesign step followed by a relax step. 14,000 decoys were produced per heme site. The spanfile defined all helical residues as membrane embedded, and starting structures were oriented in the membrane using the PPM server. Decoys were scored using the *franklin2019* score function with polar pore estimation disabled. The quality of repacked cores was assessed using the Packstat metric. The membrane topology of designed sequences was predicted using DeepTMHMM^51^. All files associated with Rosetta design and relaxation can be found at: https://github.com/BJHardy/monoheme_design.

### 4.2 Soluble protein design

The sequence of m2-4D2 was created by introducing the m4D2 mutations into the opposite ends of each helix of 4D2, effectively swapping the heme-binding and core-packing halves around. No further design was performed. The mutations from 4D2 to inverse-m1-4D2 were G20W, M23L, H37L, A41I, Y75I, G78W, M81L, H95F and A99L. A Rosetta model of m2-4D2 was produced by threading the mutated sequence onto the crystal structure of 4D2 (with reconstructed loops) using the SimpleThreadingMover followed by FastRelax implemented in RosettaScripts (see Supplementary). A predicted structure was also produced with ESMfold^45^. All files associated with Rosetta sequence threading can also be found in the GitHub repository.

### 4.3 Protein structure prediction

Sequences were subject to structure prediction with ESMfold via the ‘ESMfold_advanced’ Google Colab environment, with num_recycles set to 3 and the language models contacts option enabled: (https://colab.research.google.com/github/sokrypton/ColabFold/blob/main/beta/ESMFold_advanced.ipynb). Structure prediction with AlphaFold2 was performed via the ColabFold environment (https://colab.research.google.com/github/sokrypton/ColabFold/blob/main/AlphaFold2.ipynb#scrollTo=kOblAo-xetgx) in SingleSequence mode, with relaxation disabled and all other settings as default. Alignments of predicted structures and experimental or Rosetta models of proteins was performed using the TM-score web server (https://zhanggroup.org/TM-score/)^62^.

### 4.4 Mass spectrometry

The molecular weight of m1-CytbX and m2-CytbX holoproteins was analysed by native mass spectrometry as previously described for CytbX^43^. The molecular weight of heme-bound m2-4D2 was measured as described previously for 4D2 and m1-4D2^42^.

### 4.5 Molecular biology

Synthetic genes were ordered in expression vectors from Twist Bioscience. The m2-4D2 gene was purchased cloned between the EcoRI and NotI sites of a pET-21+ vector, with a N-terminal 6xHis tag followed by a tobacco-etch virus (TEV) cleavage site. Synthetic genes encoding 10xHis-tagged CytbX-Mono proteins, sfGFP fusions and Strep3-tagged m1-CytbX-0016 and m2-CytbX-0320 were ordered from Twist Biosciences cloned into the NdeI/XhoI sites of pET-29(b)+.

### 4.6 Protein expression and purification

Expression and purification methods of membrane proteins (m1-CytbX, m2-CytbX) and soluble proteins (m2-4D2) was as described previously^42,43^. Membrane proteins were expressed in C43 (DE3) *E. coli* and soluble proteins were expressed in T7 express *E. coli* (NEB). Affinity purification of Strep3-tagged proteins was performed using a Strep-Tactin® Superflow® high capacity FPLC column (IBA Lifesciences).

### 4.7 Protein characterisation

UV-visible (UV-Vis) absorbance spectra of purified proteins were measured from 250-750 nm in a quartz cuvette using a Cary UV-Vis spectrophotometer. The stoichiometry of ZnPPIX binding to m2-CytbX was estimated using calculated extinction coefficients of bound ferrous heme at 533 nm of 12,400 M^-1^cm^-1^ and bound ZnPPIX at 592 nm of 23,685 M^-1^cm^-1^. Fluorescence spectra of proteins containing ZnPPIX were measured from 550-750 nm in a Cary Eclipse Fluorescence Spectrophotometer (Agilent Technologies) with an excitation wavelength of 430 nm. Two prominent peaks in the fluorescence emission spectrum of m2-CytbX at 593 nm and 648 nm confirmed bound ZnPPIX^53^. Optically-transparent thin-layer electrochemistry (OTTLE) measurements of membrane and soluble proteins were performed and analysed as described previously^42,43^. OTTLE experiments were performed at pH 7.4 for membrane proteins and at pH 8.6 for water soluble proteins. Midpoint potentials for proteins designed in this study are reported as the mean potential derived from single-electron Nernst fits to three separate experiments.

### 4.8 Continuum-Electrostatics calculations

The redox potential shifts of the heme groups in 4D2, m1-4D2, m2-4D2, m1-CytbX and m2-CytbX were determined using a combination of Poisson–Boltzmann (PB) calculations and Metropolis Monte Carlo (MC) simulations as described in detail previously^63^. This method, which involves the simulation of the joint binding equilibrium of proton and electrons, uses PB calculations with the MEAD software^64^ and MC calculations with the software PETIT^65^. The (individual and pairwise) terms needed for the free energies associated with protonation/reduction changes are computed using the PB equation. Such energies are then used in the MC calculations.

The atomic charges and radii for all the atoms in the protein and heme group were taken from the GROMOS 54A7 force field^66^ and from our previous work^42^, respectively. The simulations used a temperature of 298 K and a molecular surface defined with a solvent probe radius of 1.4 Å. Dielectric constants of 80 and 20 were used for the solvent and protein^63^, respectively. The membrane was modelled as a low-dielectric slab parallel to the x-y plane using a dielectric constant value of 20. Each MC simulation comprises 10^5^ MC steps, and the acceptance/rejection of each step followed a Metropolis criterion using the previously determined PB free energies.

Predicted reduction potentials of hemes were obtained by fitting the simulated titration curves for the hemes of m1-4D2, m2-4D2, m1-CytbX and m2-CytbX to a Nernst equation describing a single electron reduction event. This approach was previously shown to perform well in predicting redox potential shifts for m1-4D2 and its mutants^42,67^.

### 4.8 BioDC Methods

The structures of 4D2, m1-4D2, m2-4D2, CytbX, m1-CytbX, and m2-CytbX were provided in PDB format to the Structure Preparation & Relaxation module of BioDC. This module interactively assists the user in preparing Atomic Model Building with Energy Refinement (AMBER) topology and coordinate files, The user can (1) select to mutate none, one, or multiple residues; (2) designate Asp, Glu, His, Tyr, and Lys residues as titratable for constant pH molecular dynamics simulations; (3) select pairs of Cys residues that should have disulphide linkages; (4) assign the reduced or oxidised state to each heme individually; and (5) decide to immerse or not the protein in an aqueous rectangular or octahedral water box with a specified thickness for the water layer, and specified numbers of Na^+^ and Cl^-^ ions. Importantly, the module automatically detects whether each heme is of the *b*- or *c*-type variety and whether each heme has His-His or His-Met axial ligands to correctly assign the appropriate force field parameters for the user-selected redox state. Only these types of hemes are currently supported by BioDC.

The parameters for His-His ligated *b*-type hemes were in part taken from Yang *et al.*^68^, and developed with the Metal Center Parameter Builder (MCPB) program^69^ of the AmberTools suite^70^ based on quantum chemical calculations using the B3LYP approximate density functional and a mixed basis set (LANL2TZ(f)) for Fe and 6-31G(d) for all second-row elements). Gaussian 16 Rev. A.03^71^ was used for the quantum chemical calculations.

The prepared structures were passed to the Energetic Estimation module of BioDC. This module interfaces with the Poisson-Boltzmann Surface Area (PBSA) program of the AmberTools suite to calculate (1) the oxidation energy of each heme while all other hemes are in the reduced state, and (2) the change in the oxidation energy of each heme due to the oxidation of another heme, while all other hemes (if there are more than two hemes in total) are in the reduced state. These quantities reflect, respectively, the influence of the electrostatic environment and heme-heme interactions on the heme redox potentials.

The interior protein static dielectric constant for the PBSA calculations was estimated from the solvent accessible surface area of the hemes^72^. Static dielectric constants of 6.776 and 6.445 were assigned to the protein interiors of 4D2 and CytbX, respectively. These values were used for the associated mono-heme variants to facilitate comparability of mono- and di-heme constructs.

4D2, m1-4D2, and m2-4D2 were modelled in an aqueous environment with a medium dielectric of 78.2. CytbX, m1-CytbX, and m2-CytbX were modelled in a membranous environment with a dielectric equal to that of the protein interior (6.445), as done previously^72^. The thickness of the implicit membrane was 30.6 (CytbX), 28.6 (m1-CytbX), and 30.4 (m2-CytbX) Å, as predicted with the PPM 3.0 server provided by the Orientation of Proteins in Membranes^73^. Beyond the membranous region of ∼30 Å centred on each CytbX-related construct, the medium dielectric was set to 78.2 for an aqueous environment.

The Energetic Estimation module of BioDC can also compute the heme-to-heme electron transfer reorganisation energy, assign a generic electronic coupling value based on the mutual orientation of the cofactors, and calculate the non-adiabatic Marcus theory electron transfer rate. None of these features were needed for the present study. Likewise, BioDC has a Redox Current Prediction module that was not needed for the present study. This module uses the estimated electron transfer rates to compute the associated redox current in the diffusive and protein-limited steady-state regimes. Results using these other features of BioDC have been preliminarily described^1^. The version 2.0 of BioDC used for the present work is available via a GitHub repository: https://github.com/Mag14011/BioDC.

### 4.9 NMR spectroscopy

Isotopic labelling of m2-4D2 was performed as previously described^74^. Briefly, the cultures were grown in LB at 37 °C, and upon reaching the OD600 of 0.8 the cells were pelleted and resuspended in cell wash solution (22 mM KH_2_PO_4_, 48 mM Na_2_HPO_4_, 8.6 mM NaCl). The cultures were then pelleted again, and the cells from 3 L of bacterial cultures were gently resuspended in 750 mL of M9 minimal media (22 mM KH_2_PO_4_, 48 mM Na_2_HPO_4_, 8.6 mM NaCl, 18 mM ^15^N-NH_4_Cl, 4 g/L glucose, 1 x Basal Medium Eagle Vitamins (100x stock; VWR/Lonza #733-1801), 2 mM MgSO_4_, 0.1 mM CaCl_2_). These were incubated at 37 °C for 1 h to aid cell recovery and subsequently induced with 1 mM IPTG. The proteins were expressed for 4 h at 37°C and purified as normal. SEC was performed at pH 6.4 (50 mM potassium phosphate, 20 mM potassium chloride).

The concentration of the NMR samples was 350 µM in 50 mM potassium phosphate, 20 mM potassium chloride (pH 6.4) with 10% D2O. 1H-15N HSQC-TROSY spectra were acquired using standard Bruker pulse programs at 25°C on a 700 MHz Bruker AVANCE *HD III* NMR spectrometer equipped with a 1.7 mm triple-resonance micro-cryoprobe. All the NMR data were processed using Topspin 3.6 (Bruker, Coventry, UK) and the spectra were visualized and using CcpNmr Analysis version 2.4.2 ^75^ (Wim et al, 2005).

## Author Contributions

**Benjamin Hardy**: Conceptualization; investigation; writing – original draft preparation; data curation. **Paulina Dubiel**: Conceptualization; investigation; writing – review & editing; data curation. **Ethan Bungay**: Investigation; writing – review & editing. **May Rudin**: Investigation. **Christopher Williams**: Investigation; resources. **Christopher Arthur**: Investigation; resources. **Matthew Guberman-Pfeffer**: Conceptualization; investigation; funding acquisition; writing – review & editing; software; methodology; data curation. **Sofia Oliveira**: Conceptualization; investigation; funding acquisition; writing – review & editing; software; methodology; data curation. **Paul Curnow**: Conceptualization; investigation; funding acquisition; writing – review & editing. **Ross Anderson**: Conceptualization; investigation; funding acquisition; writing – original draft preparation.

## Acknowledgements

This work was supported at the University of Bristol by the Biological and Biotechnological Sciences Research Council (BBW003449/1 and BB/X009831/1). We wish to thank the University of Bristol for funding a studentship for E.L.B. M.J.G.-P. wishes to acknowledge the financial support of an Office of Vice Provost Research fellowship at Baylor University and the mentorship of Bryan Shaw.

