## Supplementary information for "Delineating redox cooperativity in water-soluble and membrane multiheme cytochromes through protein design"

**Figure S1.** Structure predictions of m2-4D2 with ESMfold and AF2.

**Figure S2.** Comparison of helical packing order in predicted structures from AF2 vs. the theoretical Rosetta-modelled structure of m2-4D2.

**Figure S3.** Comparison of ESMfold and AF2 predicted structures vs the crystal structure of 4D2.

**Figure S4.** Structure predictions of m1-4D2 with ESMfold and AF2.

**Figure S5.** Overlay of ESMfold predicted structures vs. target structure of diheme and monoheme proteins.

**Figure S6**. ESMfold structure prediction of CytbX containing the nine core packing mutations from m4D2.

**Figure S7.** Score and sequence similarity analysis for the top 10 Rosetta-designed sequences for m1-CytbX and m2-CytbX.

**Figure S8.** ESMfold structure predictions of m1-CytbX and m2-CytbX.

**Figure S9.** AlphaFold2 structure predictions of m1-CytbX and m2-CytbX.

**Figure S10.** DeepTMHMM predicted transmembrane topologies of designed membrane protein sequences.

**Figure S11.** Size-exclusion chromatography elution traces.

**Figure S12.** Spectral evidence for ZnPPIX binding to m2-CytbX

**Figure S13.** Native mass spectrometry of m1-CytbX, m2-CytbX and m2-4D2.

**Figure S14.** Expected and observed masses from native mass spectrometry of m1-CytbX, m2-CytbX and m2-4D2.

**Figure S15**. NMR spectroscopy of m2-4D2.

**Figure S16.** Electrostatic maps and isosurfaces.

**Figure S17**. Amino acid and DNA sequences.

**Table S1.** ESMfold and AF2 Structure prediction metrics for designed heme proteins.

**Table S2.** Structural deviations of ESMfold predicted structures vs target structures.

**Table S3.** Expected and observed masses from native MS.

**Table S4.** Experimental and calculated reduction potentials of hemes.

**Table S5.** Calculated oxidation energies from BioDC.

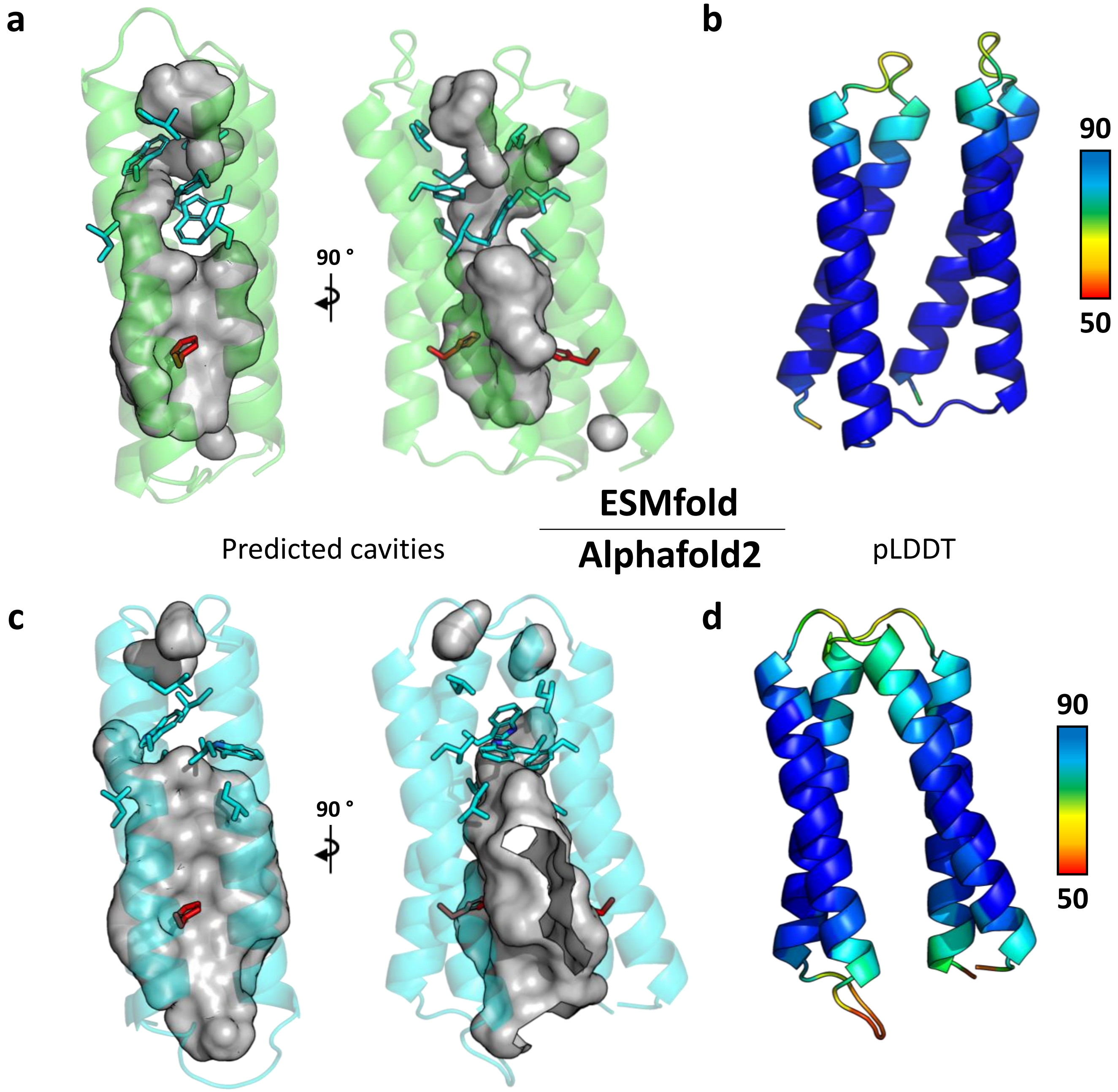


Figure S1. Structure predictions of m2-4D2 from ESMfold (top) and AlphaFold2 (bottom). (a,c) Depictions of predicted cavities, highlighting heme binding and core packing regions and (b, d) predicted structure coloured by confidence (pLDDT), as a rainbow from red to blue scaled from 50 to 90. Cavities were generated with PyMol. Histidine residues and core packing residues are shown as sticks, coloured red and cyan respectively consistent with Figure 1.

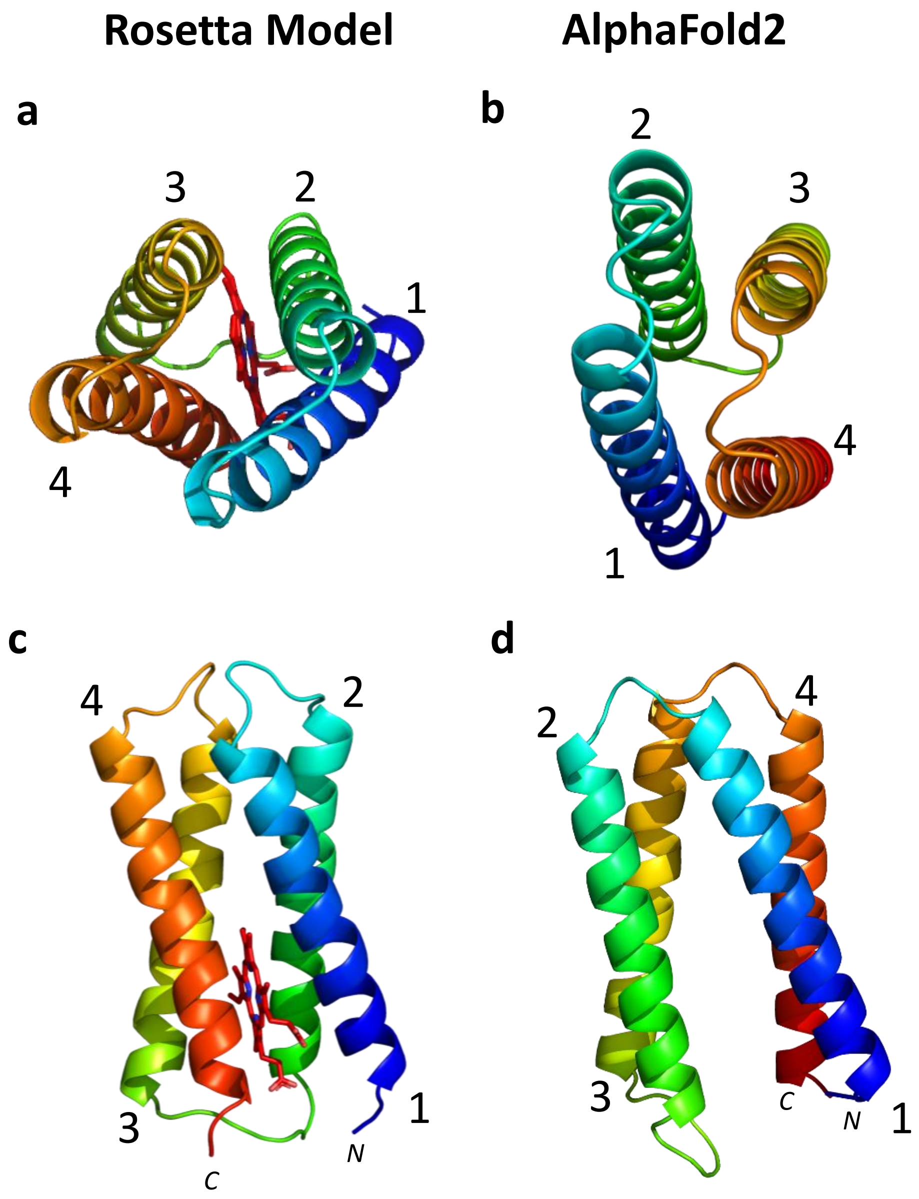


Figure S2. Comparison of helical packing order in predicted structures from AF2 vs. the theoretical Rosetta-modelled structure of m2-4D2. (a-b) Top views and (c-d) side view of helical packing. Models are coloured from N to C terminus as a rainbow from blue to red. The number of each helix in the protein is labelled. AlphaFold2 predicts a mirrored packing order to the theoretical structure.

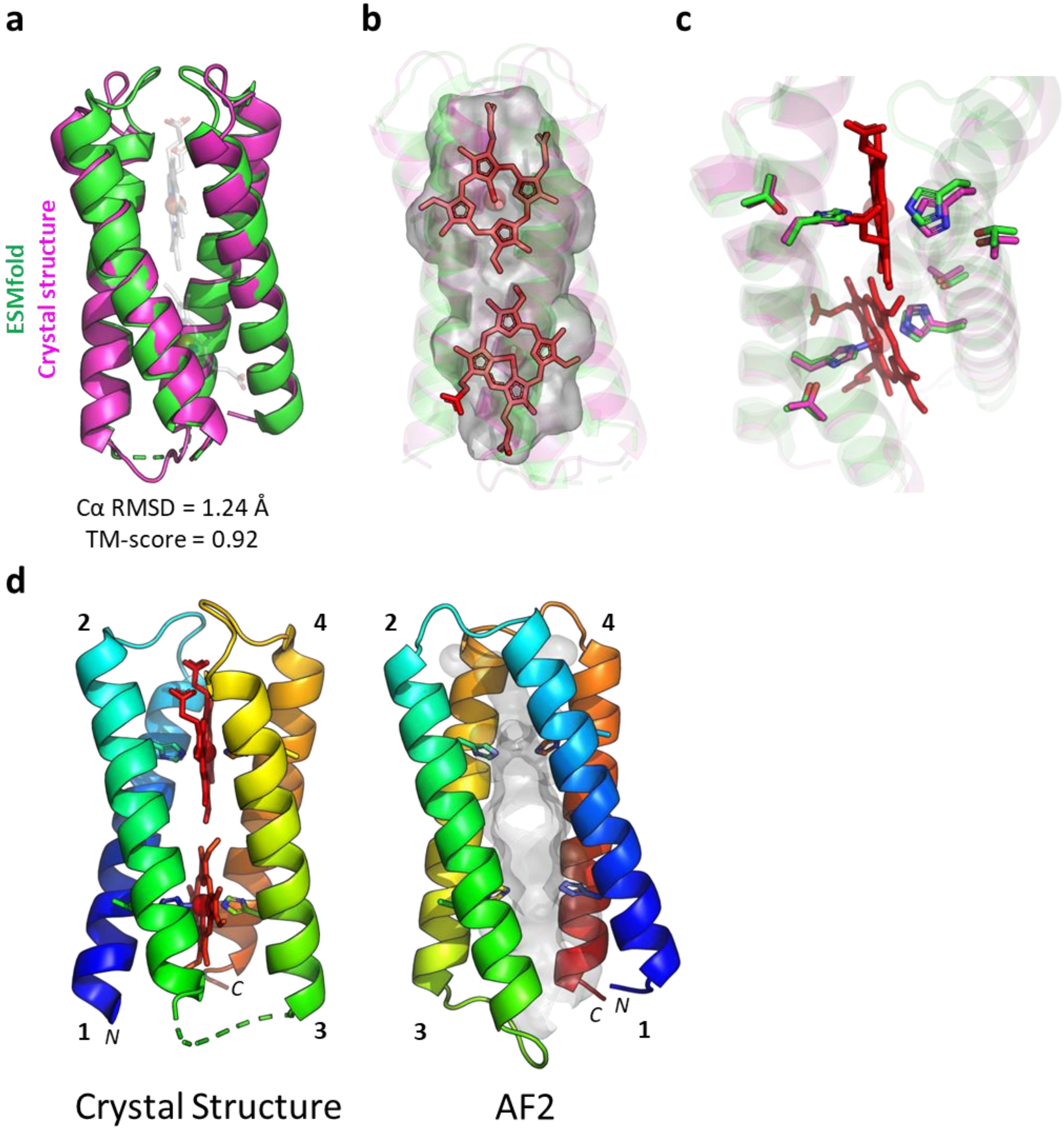


Figure S3. Comparison of ESMfold and AF2 predicted structures vs the crystal structure of 4D2 (PDB ID: 7AH0). (a) Alignment of ESMfold model (green) vs 4D2 crystal structure (magenta). (b) Heme-shaped cavities within the ESMfold predicted structure, showing the hemes of 4D2. (c) The heme-coordinating histidine-threonine residue pairs align almost exactly between the 4D2 crystal structure and ESMfold model. (d) The crystal structure and AF2 model coloured as a rainbow from N to C terminus, with hemes and internal cavities shown. The helix numbers are labelled, showing that AF2 packs the 4D2 helical bundle in a mirrored image to the crystal structure.

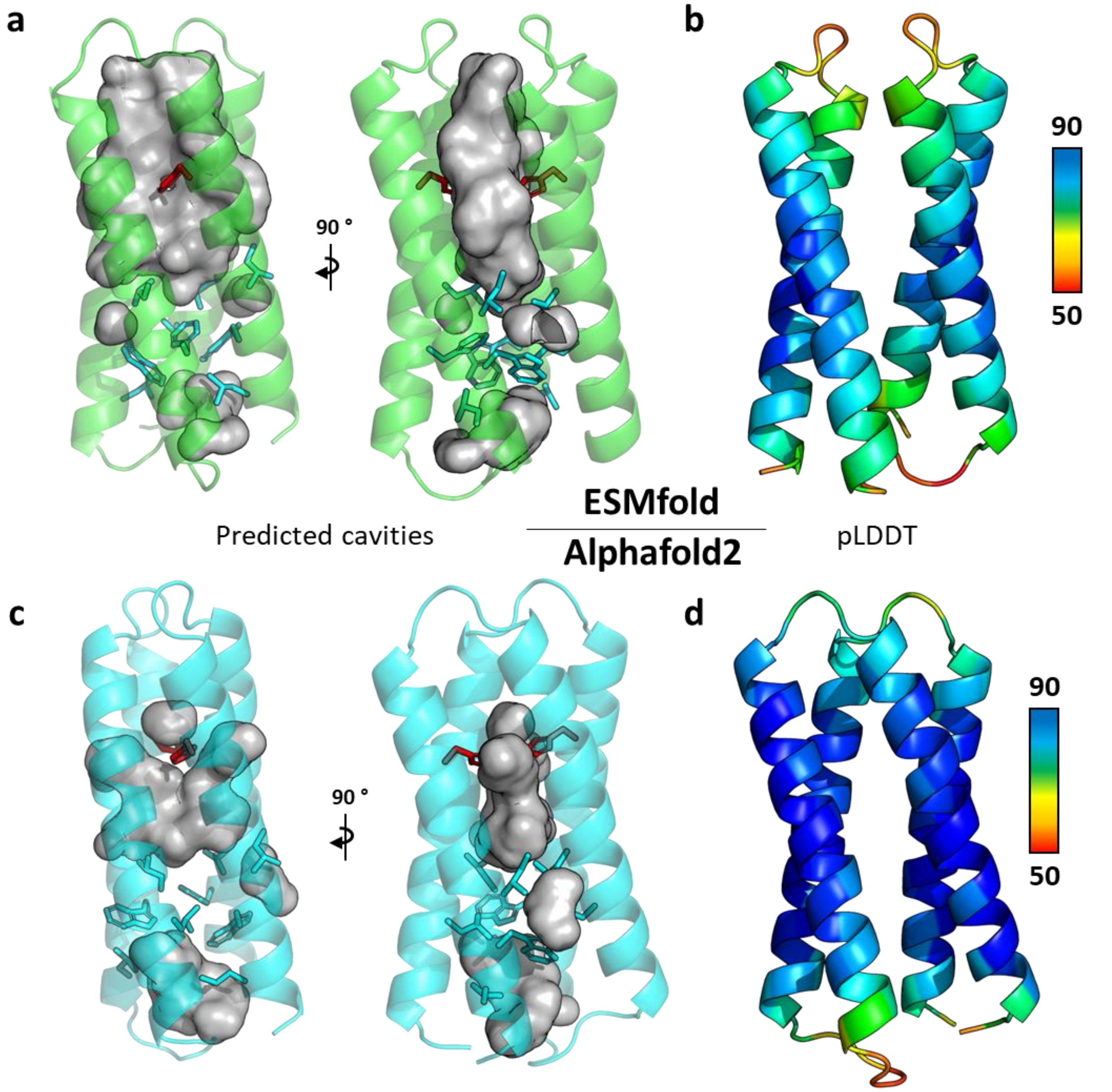


Figure S4. Structure predictions of m1-4D2 from ESMfold (top) and AlphaFold2 (bottom). (**a**,**c**) Depictions of predicted cavities, highlighting heme binding and core packing regions and (b, d) predicted structures coloured by confidence (pLDDT), as a rainbow from red to blue scaled from 50 to 90. Cavities were visualised with PyMol. Histidine residues and core packing residues are shown as sticks, coloured red and cyan respectively consistent with Figure 1.

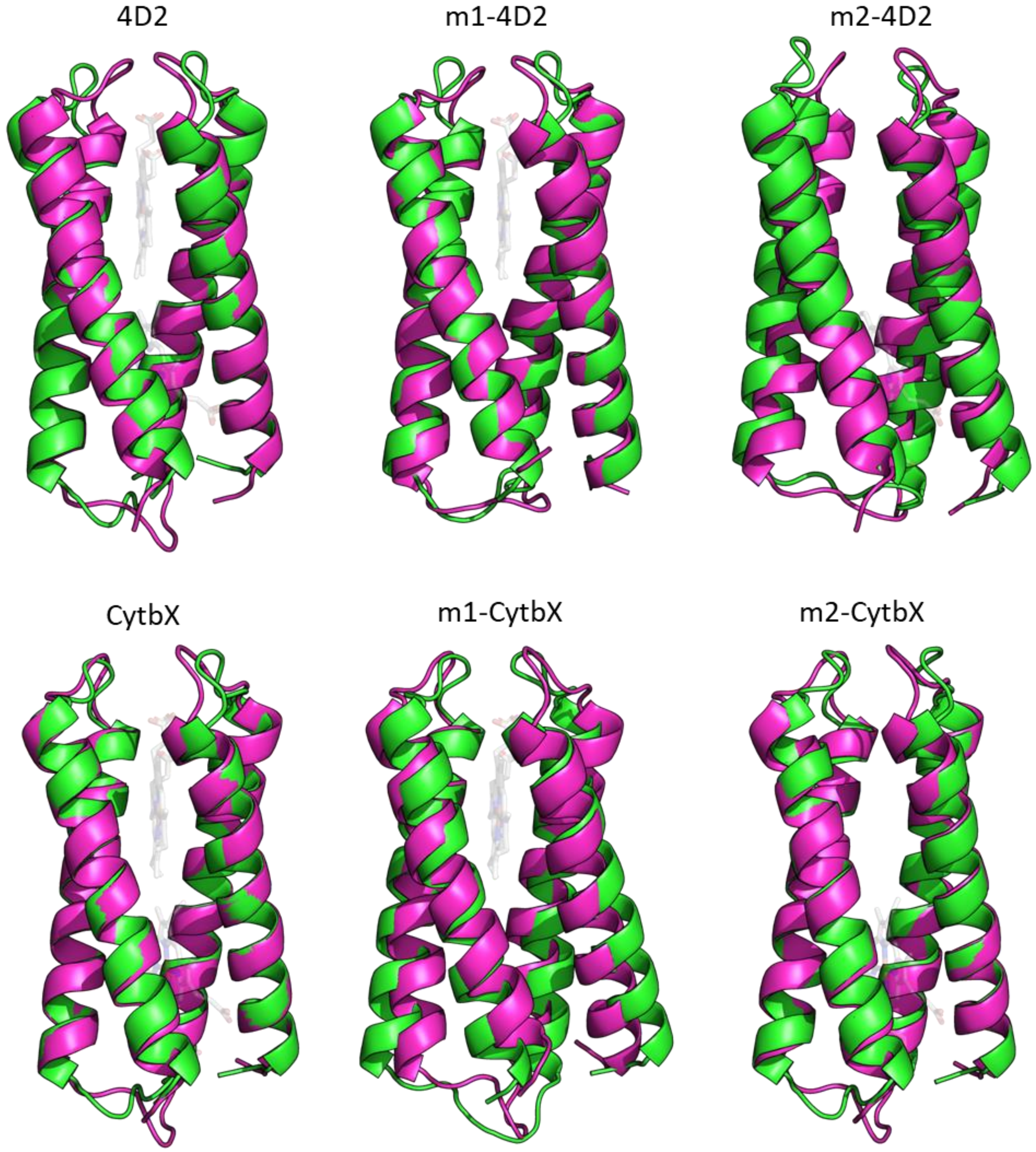


Figure S5. Overlay of ESMfold predicted structures vs. crystal structure (4D2) or modelled theoretical structures of the designed heme proteins. Green: ESMfold structure, magenta: Crystal structure (4D2) or Rosetta structure. Hemes are shown as semi-transparent sticks and are absent in predicted structures. For RMSD and TM metrics for structural alignments see supplementary table 2.

| Protein | ESMfold | | AlphaFold2 | |
|---|---|---|---|---|
| | pTM | Mean pLDDT | pTM | Mean pLDDT |
| CytbX | 0.835 | 88.6 | 0.60 | 79.5 |
| m1-CytbX (0016) | 0.852 | 89.7 | 0.73 | 86.0 |
| m1-CytbX (0151) | 0.823 | 86.8 | 0.81 | 90.8 |
| m2-CytbX (0320) | 0.836 | 86.6 | 0.52 | 77.7 |
| m2-CytbX (0152) | 0.811 | 86.6 | 0.55 | 78.3 |
| CytbX m4D2 copy | 0.811 | 86.3 | 0.64 | 83.1 |
| 4D2 | 0.814 | 85.3 | 0.56 | 79.9 |
| m1-4D2 | 0.665 | 77.7 | 0.66 | 82.8 |
| m2-4D2 | 0.810 | 87.0 | 0.57 | 82.3 |

Table S1. Structure prediction metrics for designed heme proteins. ESMfold was run using default settings with 3 recycles. AlphaFold2 was run in single sequence mode with 6 recycles, and metrics are reported for the top ranked model. Mean pLDDT reports the average pLDDT across all residues.

| Protein | ESMfold | | AlphaFold2 | |
|---|---|---|---|---|
| | Ca RMSD (Å) | TM | Ca RMSD (Å) | TM |
| 4D2* | 1.24 | 0.92 | 11.41 | 0.42 |
| m1-4D2 | 1.38 | 0.93 | 11.04 | 0.46 |
| m2-4D2 | 2.63 | 0.73 | 11.58 | 0.40 |
| CytbX | 1.29 | 0.93 | 10.35 | 0.43 |
| CytbX (m4D2 mutations) | 2.22 | 0.80 | 10.33 | 0.43 |
| m1-CytbX | 2.14 | 0.82 | 10.46 | 0.42 |
| m2-CytbX | 1.45 | 0.91 | 10.63 | 0.46 |

Table S2. Structural deviations of ESMfold predicted structures vs target structures for the final six soluble and membrane heme proteins. *Calculated vs the crystal structure of 4D2 (PDB ID: 7AH0). All others were calculated vs. the Rosetta model of each protein. RMSD and TM were calculated using the TM-score web server. The top ranked model was used from the AF2 output.

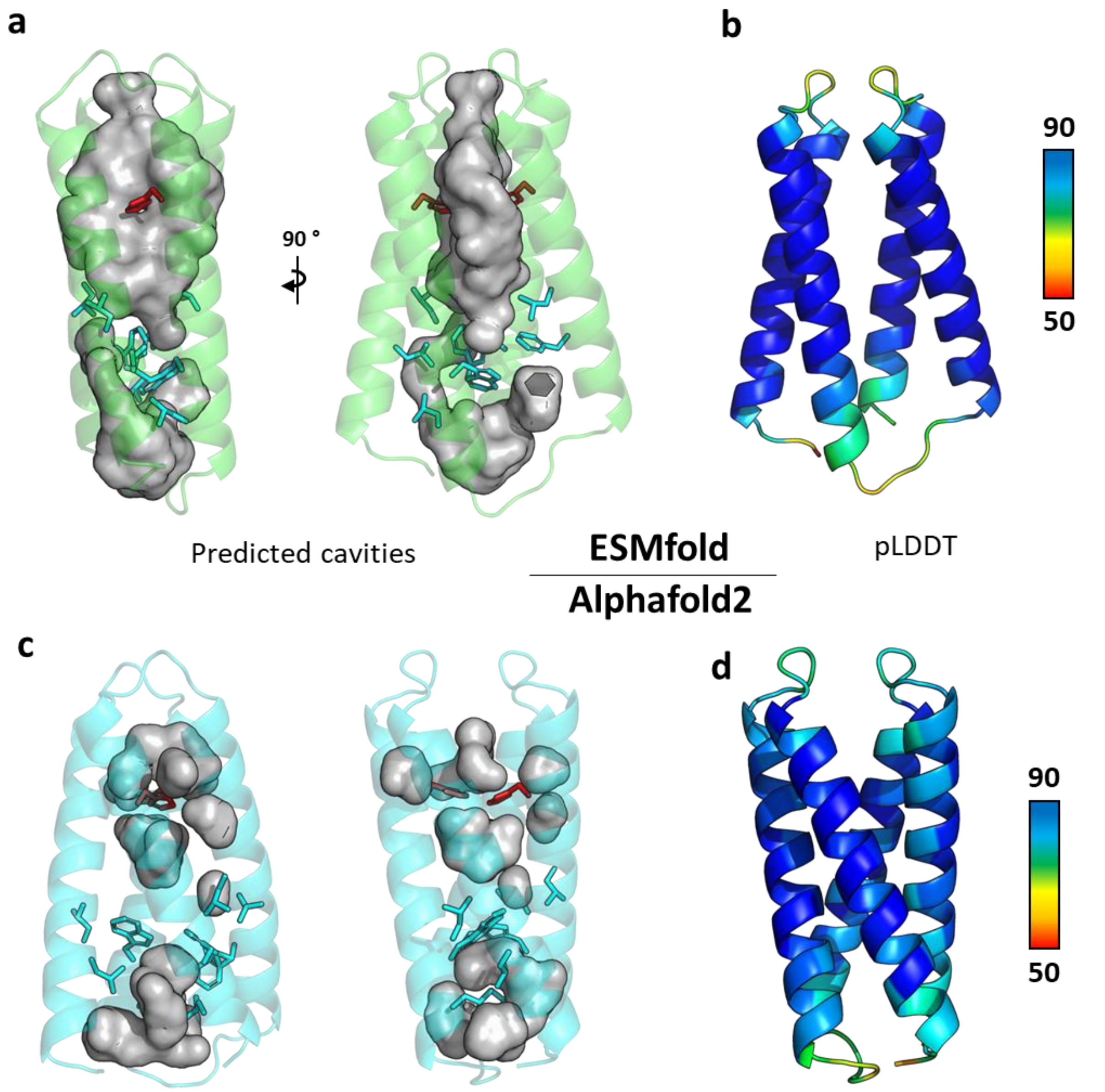


Figure S6. ESMfold structure prediction of CytbX containing the nine core packing mutations from m4D2. (a) Representation of the cavities within the ESMfold predicted structure. A heme-shaped hole can be seen in the binding site with histidine side chains pointing towards a central hole in the cavity reminiscent of bis-histidine heme ligation. A cavity can also be seen in the other end of the bundle showing that the m1-4D2 mutations are not predicted to form a well packed core. Red: histidines, cyan: m1-4D2 mutations. (b) ESMfold model coloured by pLDDT (low to high, red to blue). Respective (c) cavity and (d) pLDDT representations of the AlphaFold2 predicted structure. AlphaFold2 predicts a lower-confidence, incorrectly packed structure.

**a**

**m1-CytbX**

| Rosetta score (REU) | Cα RMSD vs. CytbX (Å) | pStat | Rosetta decoy name |
|---|---|---|---|
| **-472.843** | 0.787 | 0.500 | 10914940CytbX_Rosetta_Monoheme23_flexV1KIH_0151* |
| **-472.819** | 0.771 | 0.529 | 10914953CytbX_Rosetta_Monoheme26_flexV1KIH_0049 |
| **-472.697** | 0.775 | 0.557 | 10914885CytbX_Rosetta_Monoheme10_flexV1KIH_0400 |
| **-472.539** | 0.675 | 0.574 | 10914932CytbX_Rosetta_Monoheme19_flexV1KIH_0371 |
| **-472.257** | 0.807 | 0.550 | 10914940CytbX_Rosetta_Monoheme23_flexV1KIH_0065 |
| **-471.664** | 0.764 | 0.545 | 10914938CytbX_Rosetta_Monoheme21_flexV1KIH_0492 |
| **-471.618** | 0.785 | 0.516 | 10914881CytbX_Rosetta_Monoheme6_flexV1KIH_0014 |
| **-471.473** | 1.017 | **0.606** | 10914881CytbX_Rosetta_Monoheme6_flexV1KIH_0016** |
| **-471.343** | 0.677 | 0.547 | 10914887CytbX_Rosetta_Monoheme12_flexV1KIH_0274 |
| **-471.325** | 0.677 | 0.570 | 10914879CytbX_Rosetta_Monoheme4_flexV1KIH_0433 |

**b**

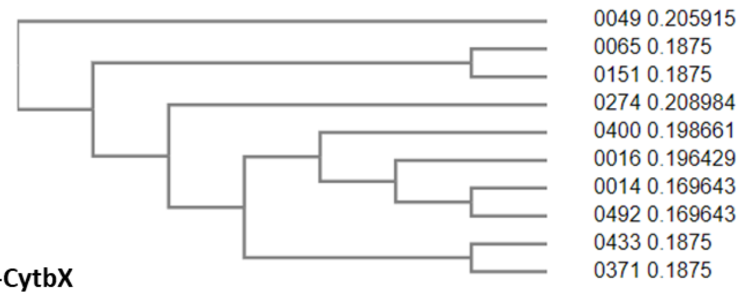


**c**

**m2-CytbX**

| Rosetta score (REU) | Cα RMSD vs. CytbX (Å) | pStat | Rosetta decoy name |
|---|---|---|---|
| **-477.497** | 1.023 | 0.587 | 10918652CytbX_Rosetta_Monoheme27_flexV1KIH_0152* |
| **-477.045** | 1.065 | 0.546 | 10938018CytbX_Rosetta_Monoheme213_flexV1KIH_0256 |
| **-475.719** | 0.960 | 0.597 | 10936424CytbX_Rosetta_Monoheme228_flexV1KIH_0114 |
| **-475.660** | 0.960 | 0.614 | 10938017CytbX_Rosetta_Monoheme212_flexV1KIH_0493 |
| **-475.584** | 0.953 | 0.615 | 10938032CytbX_Rosetta_Monoheme227_flexV1KIH_0280 |
| **-475.374** | 0.959 | **0.630** | 10938027CytbX_Rosetta_Monoheme222_flexV1KIH_0320** |
| **-475.136** | 1.020 | 0.594 | 10938018CytbX_Rosetta_Monoheme213_flexV1KIH_0235 |
| **-474.967** | 0.936 | 0.583 | 10936424CytbX_Rosetta_Monoheme228_flexV1KIH_0366 |
| **-474.814** | 0.950 | 0.613 | 10938016CytbX_Rosetta_Monoheme211_flexV1KIH_0155 |
| **-474.723** | 1.061 | 0.560 | 10938008CytbX_Rosetta_Monoheme23_flexV1KIH_0325 |

**d**

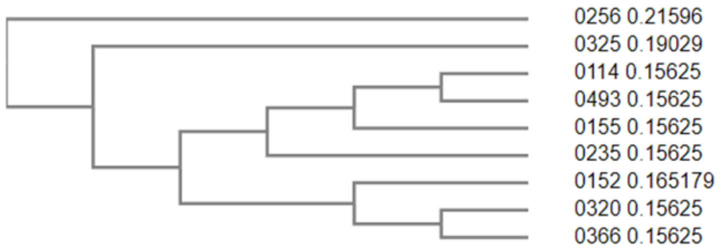


Figure S7. Score and sequence similarity analysis for the top 10 Rosetta-designed sequences for m1-CytbX and m2-CytbX. (a, c) Rosetta score, RMSD and Packstat scores for the top 10 Mono1 and Mono2 sequences, ranked by Rosetta score. Sequences were aligned in ClustalOmega, and are coloured by unique sequence identity, as identified by the guide tree. (b, d) Guide trees (as cladograms) generated by ClustalOmega showing sequence similarity of the top decoys. *Highest ranking decoy. **Highest Packstat within the top 10 decoys.

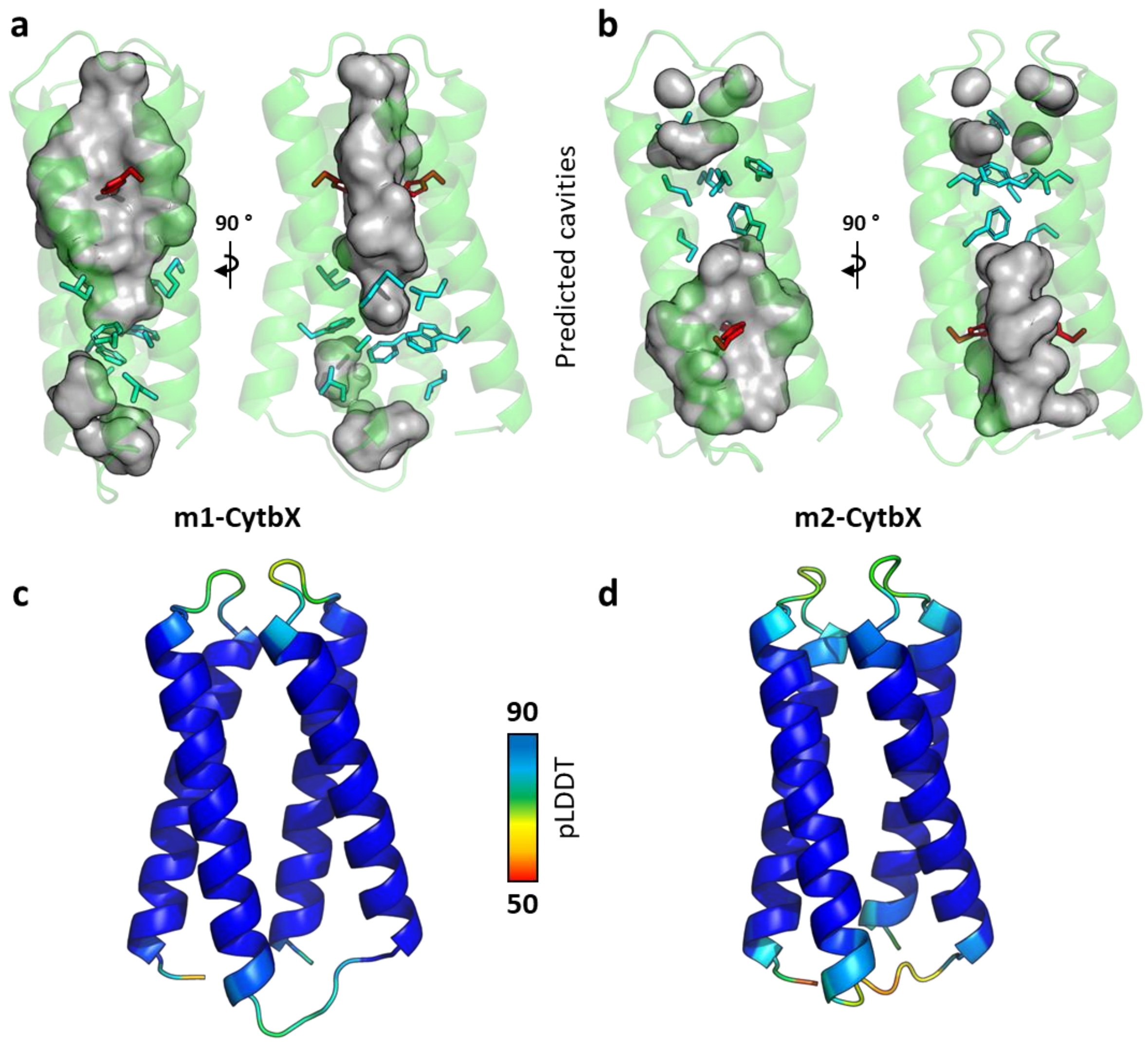


Figure S8. ESMfold structure predictions of m1-CytbX and m2-CytbX. (a-b) Depictions of predicted heme-shaped cavities within both proteins, and compact designed cores. Cavities were as calculated by PyMol. Histidine residues are shown as red sticks and designed core packing residues are shown as cyan sticks. (c-d) Predicted models coloured by confidence (pLDDT), as a rainbow from red to blue scaled from 50 to 90.

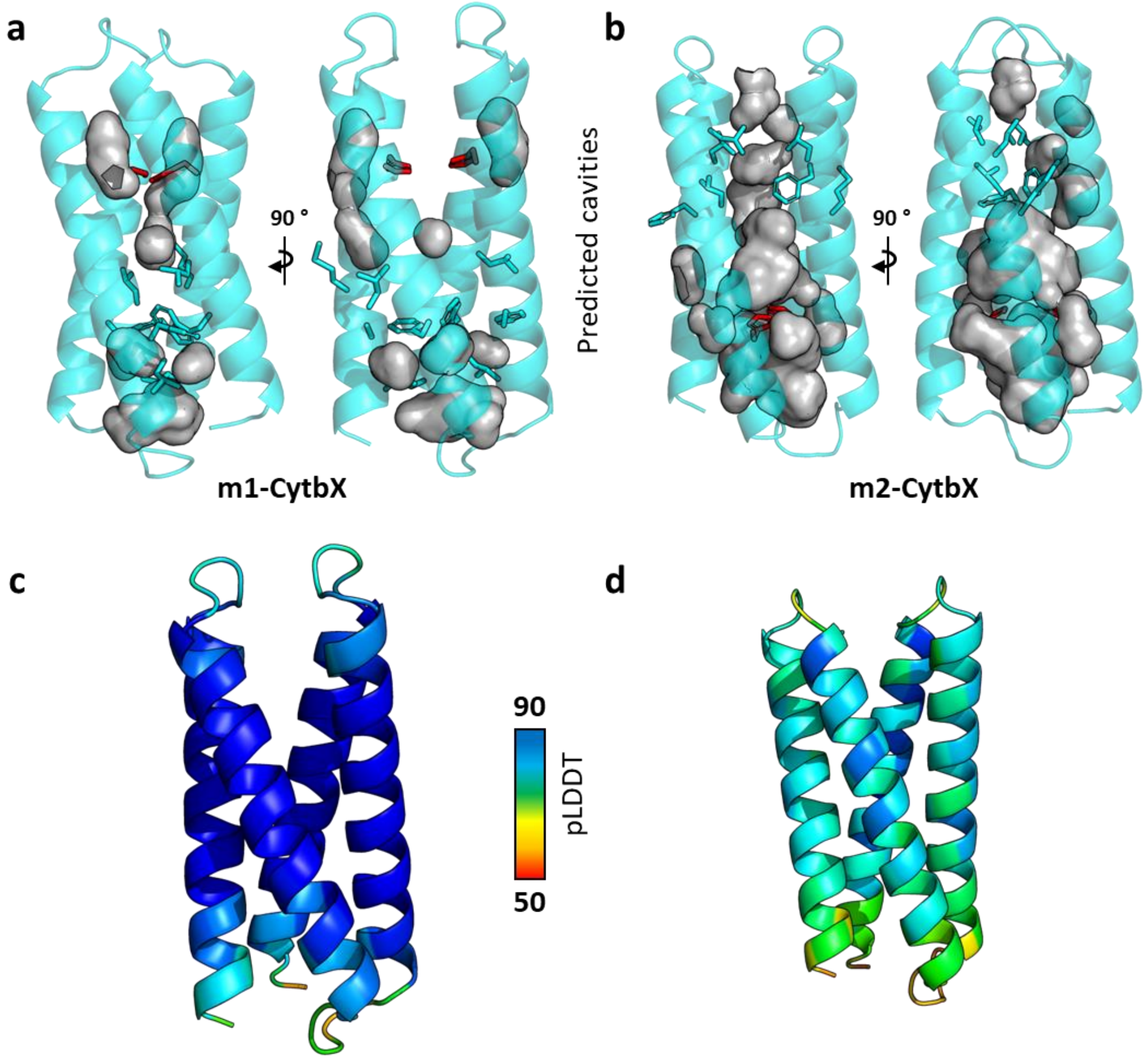


Figure S9. AlphaFold2 structure predictions of m1-CytbX and m2-CytbX. (a-b) Depictions of predicted heme-shaped cavities within both proteins, and compact designed cores. Cavities were as calculated by PyMol. Histidine residues are shown as red sticks and designed core packing residues are shown as cyan sticks. (c-d) Predicted models coloured by confidence (pLDDT), as a rainbow from red to blue scaled from 50 to 90. AlphaFold2 was run in Single Sequence mode.

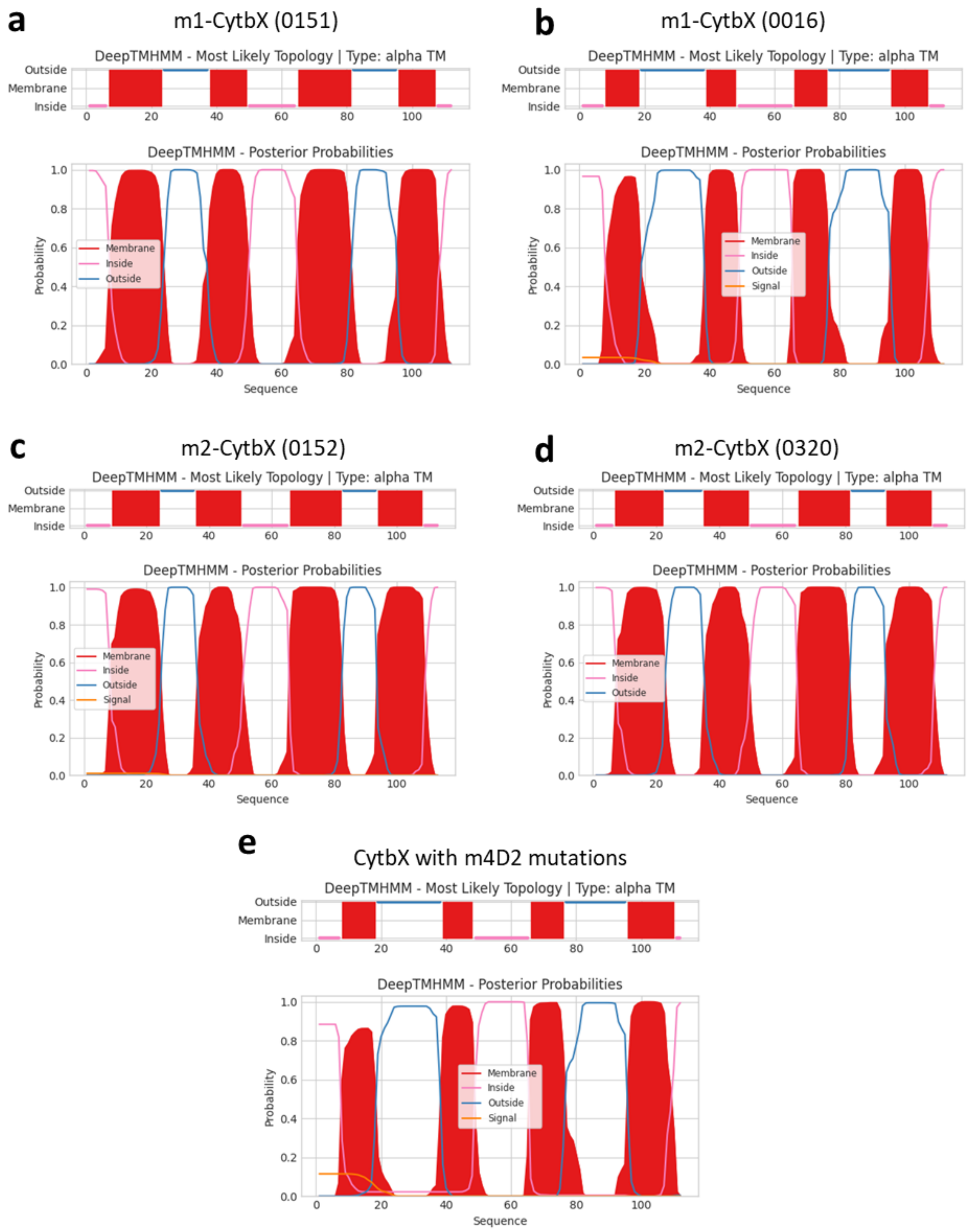


Figure S10. DeepTMHMM predicted transmembrane topologies of designed membrane protein sequences. (a-d) The four selected Rosetta-designed CytbX-Mono sequences and (e) the sequence of CytbX containing the m4D2 core mutations.

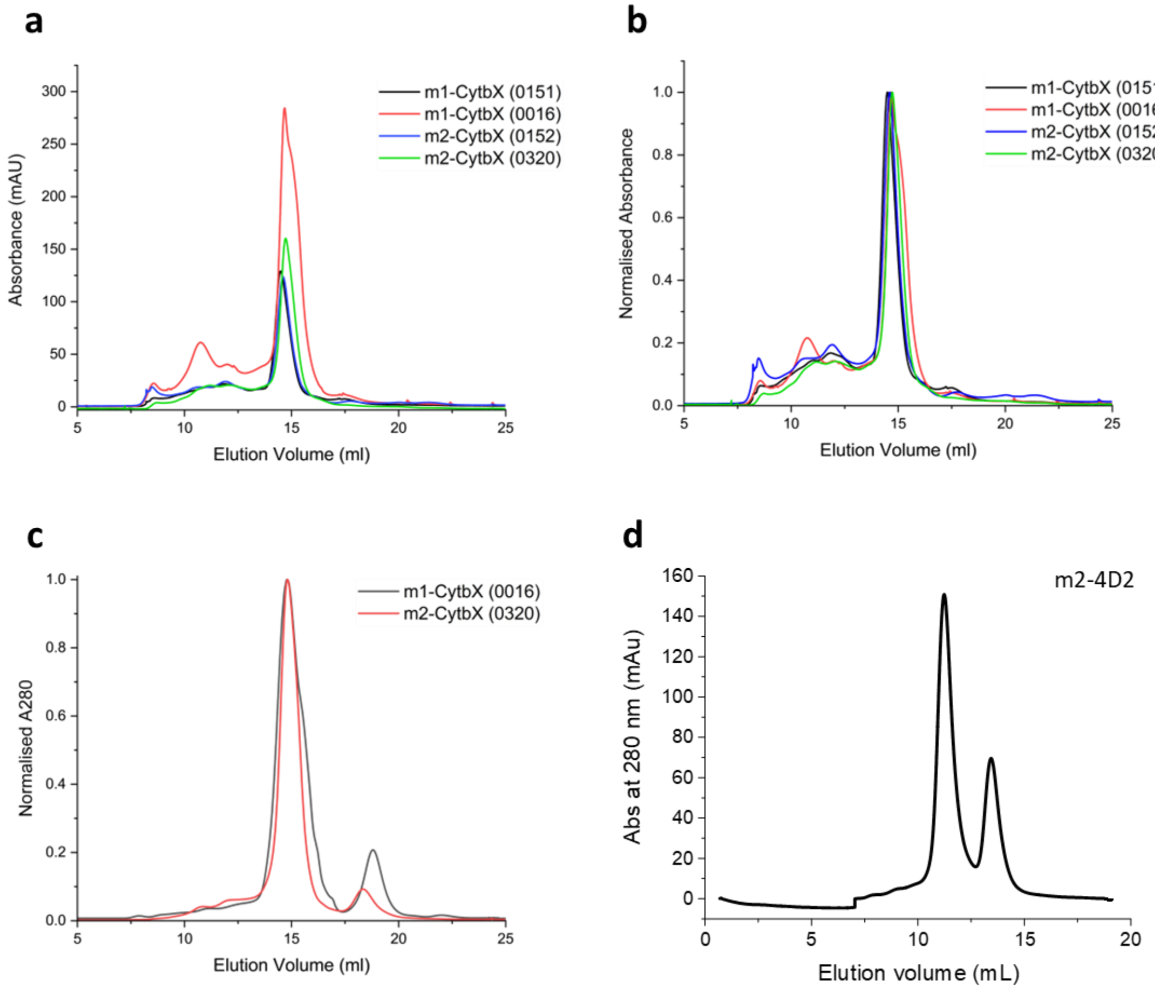


Figure S11. Size-exclusion chromatography elution traces of designed hemoproteins. (A) Raw and (b) normalised elution traces for the four monoheme CytbX sequences purified with Cymal-5, expressed with 10xHis tags. As can be seen, m1-CytbX 0016 and m2-CytbX 0320 are the best expressing of each pair and are therefore discussed in detail in the accompanying paper. (c) Normalised elution traces of m1-CytbX 0016 and m2-CytbX 0320 with triple-Strep(II) tags removed. (d) Raw elution trace for m2-4D2 after heme loading. Absorbance was measured at 280 nm.

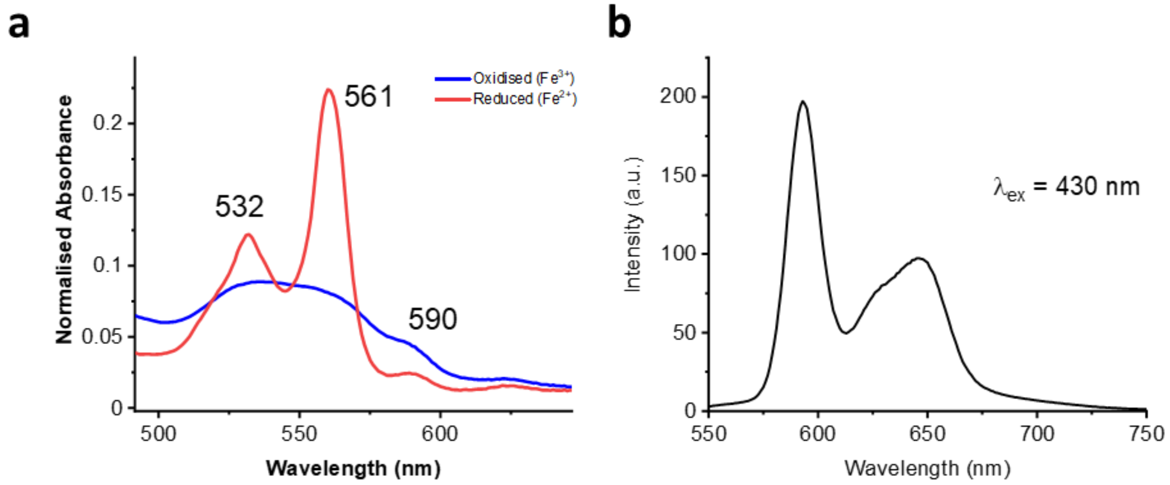


Figure S12. Spectral evidence for ZnPPIX binding to m2-CytbX. (a) UV-Vis absorbance spectrum of purified m2-CytbX with purification tags cleaved, zoomed into the region of the Q-bands. The peak at 590 nm is suggestive of ZnPPIX binding, whilst the 532 nm and 561 nm peaks are characteristic of heme *b.* (b) Fluorescence spectrum of purified m2-CytbX, excited at 430 nm. Peaks at 593 nm and 647 nm are characteristic of ZnPPIX. Slit width = 10 nm. Spectra are shown for the protein with the triple-StrepII tag removed.

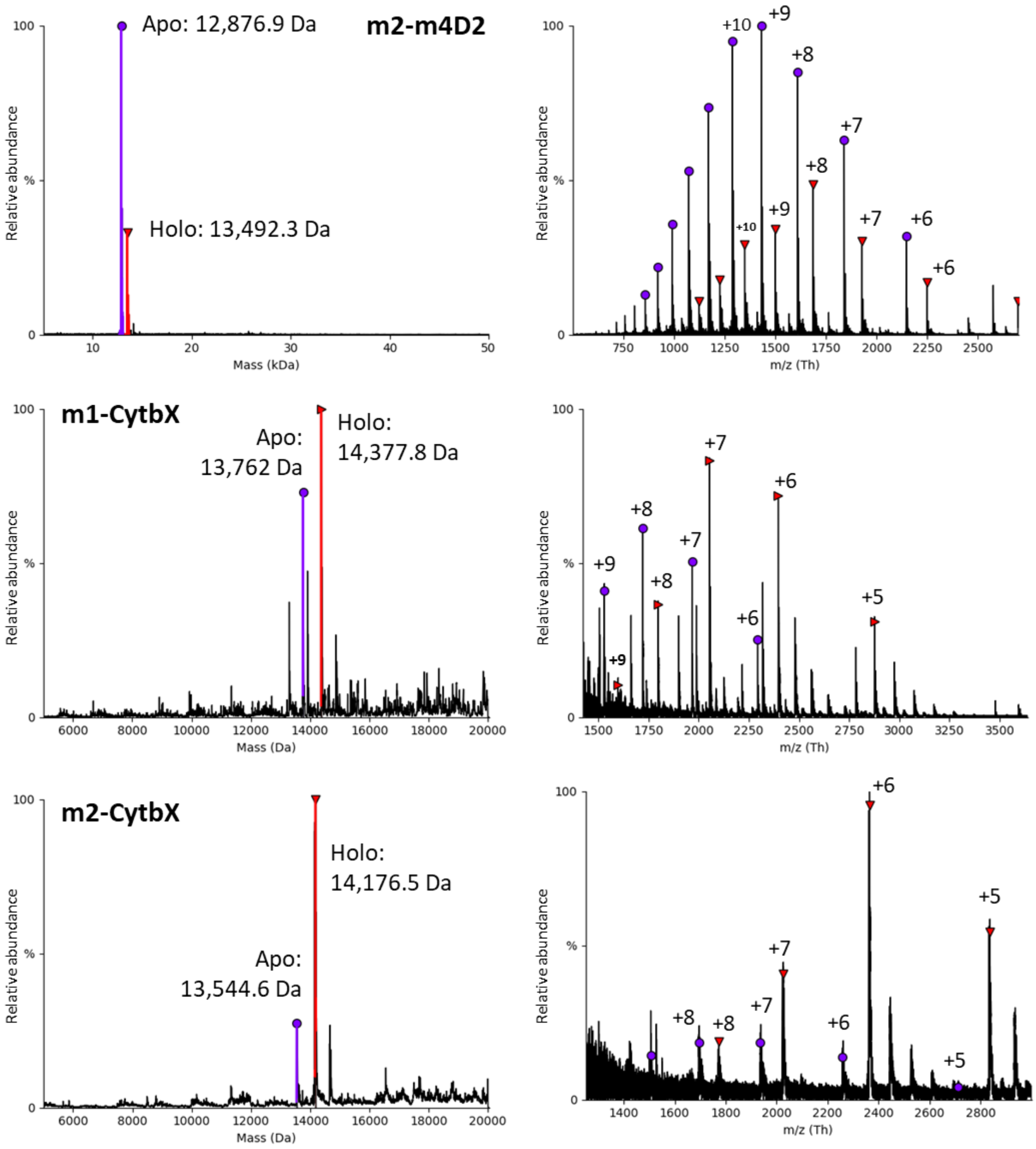


Figure S13. Native mass spectrometry of the three novel monoheme proteins. Deconvoluted (left) and raw (right) data are shown for purified proteins with purification tags removed. Purple peaks and circles represent apo-protein species and red peaks and triangles represent protein bound to a single heme (holo). Major charge state series are labelled on the m/z plots. Deconvolution of raw data was performed using UniDec, sampling masses at every 0.1 Da.

| Protein | Expected Mass Apo (Da) | Observed Mass Apo (Da) | Expected Mass Holo (Da) | Observed Mass Holo (Da) | Apo vs Holo observed mass shift (Da) |
|---|---|---|---|---|---|
| m1-CytbX | 13,735.02 | 13,762.0 | 14,351.52 | 14,377.8 | 615.8 |
| m2-CytbX | 13,513.83 | 13,544.6 | 14,130.33 | 14,176.5 | 631.9 |
| m2-4D2 | 12,874.49 | 12,876.9 | 13,490.98 | 13,492.3 | 615.4 |

Table S3. Expected and observed masses from native mass spectrometry of the three designed monoheme proteins. Expected molecular weights are reported for all proteins with purification tags removed.

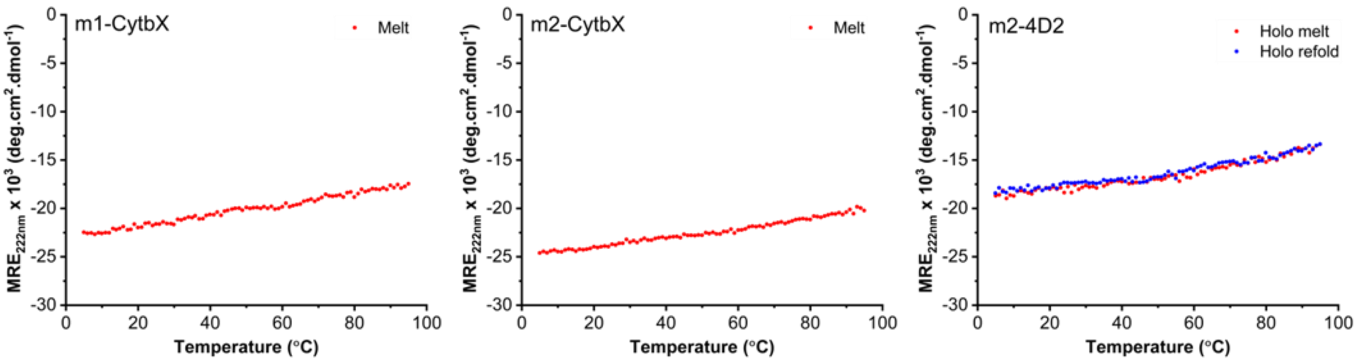


Figure S14. Thermal unfolding traces of m1-CytbX, m2-CytbX and m2-4D2. The mean residue ellipticity at 222 nm as a proxy for helicity is plotted vs temperature. All proteins show a roughly linear decrease in helicity with no major unfolding events. For m2-4D2 the trace when cooled from 95 °C to 5 °C is also overlayed in blue.

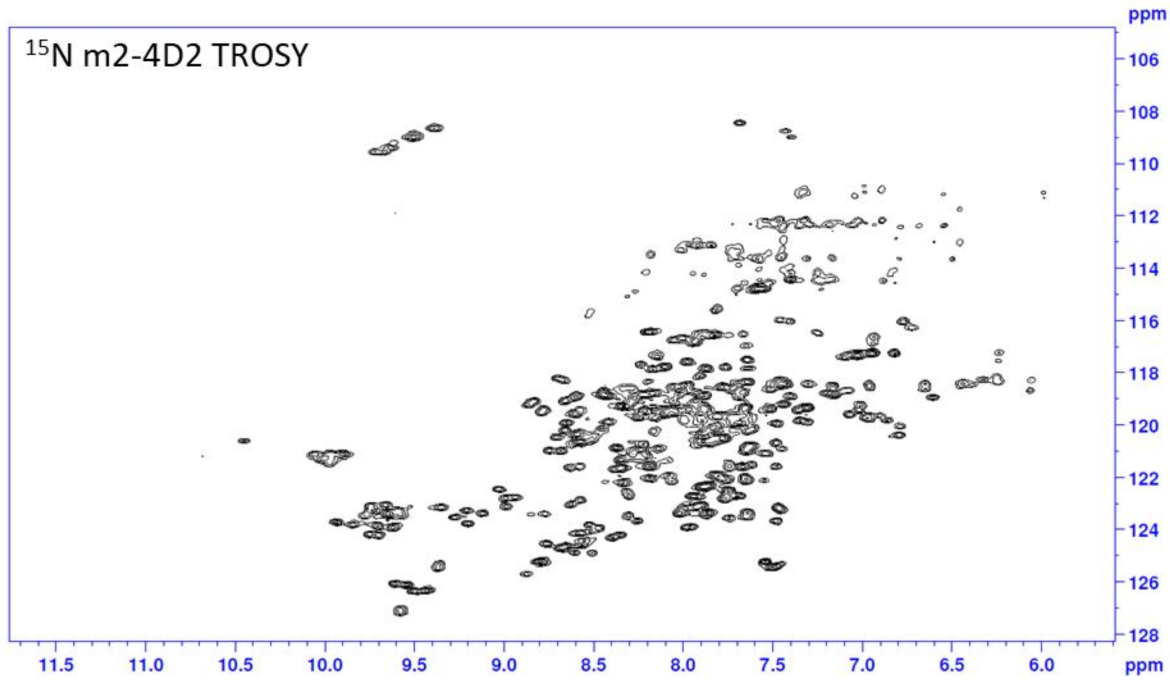


Figure S15. Nuclear magnetic resonance (NMR) spectroscopy of m2-4D2. TROSY spectrum of $^{15}$N-labelled m2-4D2 (pH 6.4, 298 K, 10% $D_2O$).

| Protein | Measured $E_m$ (mV) | Observed $\Delta E_m$ (mV) | Predicted $\Delta E_m$ (mV) (PB-MC) | Predicted $\Delta E_m$ (mV) (BioDC) |
|---|---|---|---|---|
| 4D2 (Heme 1) | -104† | 0 | n/a | 0 |
| 4D2 (Heme 2) | -167† | -63 | n/a | -51 |
| m1-4D2 | -117 | -13 (0)* | 0 | 16 (0)* |
| m2-4D2 | -125 ± 2 | -19 (-6)* | -5 | 18 (+2)* |
| CytbX (Heme 2) | -10 | 0 | n/a | 0 |
| CytbX (Heme 1) | -121 | -111 | n/a | -126 |
| m1-CytbX | -66 ± 2 | -46 (0)* | 0 | -35 (0)* |
| m2-CytbX | -82 ± 1 | -62 (-16)* | +2 | -65 (-30)* |

Table S4. Comparison of experimental and calculated reduction potentials of hemes. $\Delta E_m$ values are expressed vs. standard hydrogen electrode (SHE) relative to Heme 1 of 4D2 or Heme 2 of CytbX for soluble and membrane proteins respectively. Values in brackets (*) are expressed relative to m1-4D2 or m1-CytbX respectively. †Previously reported values.

| Protein | | $\epsilon_{\mathrm{int}}$ | $\epsilon_{\mathrm{mem}}$ | $\epsilon_{ext}$ | Charge Set | $\Delta E_{\mathrm{ox}}$ (eV) |
|---|---|---|---|---|---|---|
| 4D2 | | | | | | |
| | Site #1 | 6.776 | | 78.2 | RESP | -0.369 |
| | Site #2 | 6.776 | | 78.2 | RESP | -0.420 |
| | Site #1 | 6.776 | | 78.2 | CM5 | 0.006 |
| | Site #2 | 6.776 | | 78.2 | CM5 | -0.044 |
| m1-4D2 | | | | | | |
| | Site #1 | 6.776 | | 78.2 | RESP | -0.353 |
| | Site #1 | 6.776 | | 78.2 | CM5 | 0.018 |
| m2-4D2 | | | | | | |
| | Site #1 | 6.776 | | 78.2 | RESP | -0.351 |
| | Site #1 | 6.776 | | 78.2 | CM5 | 0.018 |
| CytbX | | | | | | |
| | Site #1 | 6.445 | | 78.2 | RESP | -0.307 |
| | Site #2 | 6.445 | | 78.2 | RESP | -0.233 |
| | Site #1 | 6.445 | 6.445 | 78.2 | RESP | -0.325 |
| | Site #2 | 6.445 | 6.445 | 78.2 | RESP | -0.199 |
| | Site #1 | 6.445 | 6.445 | 78.2 | CM5 | 0.002 |
| | Site #2 | 6.445 | 6.445 | 78.2 | CM5 | 0.130 |
| m1-CytbX | | | | | | |
| | Site #1 | 6.445 | 6.445 | 78.2 | RESP | -0.225 |
| | Site #1 | 6.445 | 6.445 | 78.2 | CM5 | 0.097 |
| m2-CytbX | | | | | | |
| | Site #1 | 6.445 | 6.445 | 78.2 | RESP | -0.262 |
| | Site #1 | 6.445 | 6.445 | 78.2 | CM5 | 0.065 |

Table S5. Oxidation energies computed ~~with~~ using BioDC with RESP and CM5 charge-fitting schemes for the heme group in the oxidised and reduced states. $\epsilon_{\mathrm{int}}$, $\epsilon_{\mathrm{mem}}$, and $\epsilon_{\mathrm{ext}}$ are respectively the dielectric constants assigned to the protein interior, implicit membrane slab if present, and exterior solvent.

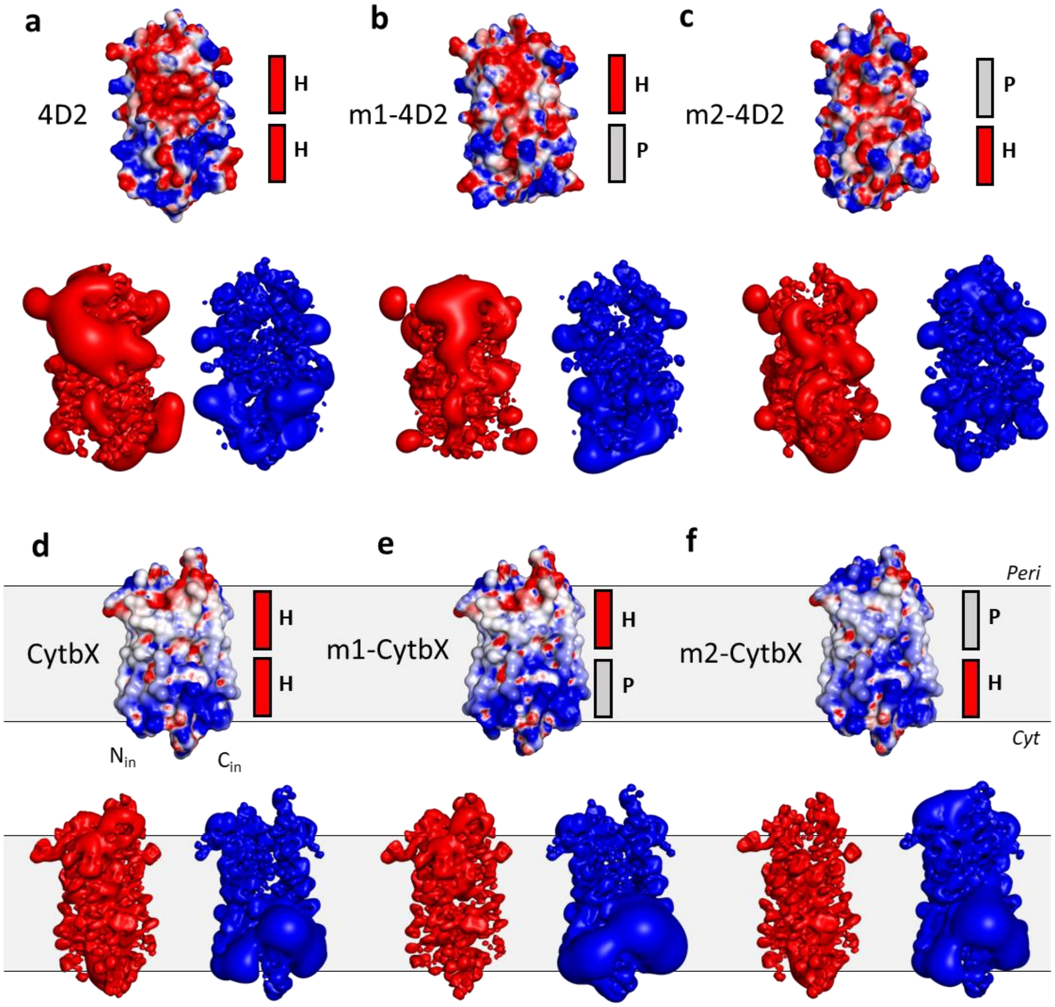


Figure S16. Electrostatic maps and isosurfaces of the two diheme and four monoheme proteins. For each protein, top: electrostatic surface map, bottom: negative (red) and positive (blue) charge isosurfaces. Red and grey boxes beside the electrostatic maps denote the positions of heme-binding (**H**) and packing (**P**) modules respectively. In all cases, proteins are positioned with N and C termini facing downwards. The position of the membrane is approximate.

Figure S17. Amino acid and DNA sequences.

All water-soluble constructs have an upstream **His tag**, **V5 epitope** **and a** **TEV cleavage site**:
**Amino acid** (AA) sequence: MHHHHHHGKPIPNPLLGLDSTENLYFQ
**DNA sequence:** ATGCATCATCACCATCACCATGGTAAGCCTATCCCTAACCCTCTCCTCG
GTCTCGATTCTACGGAAAACCTGTATTTTCAG

**>Inv-m4D2**
**Vector:** pET-21(+)
**AA sequence** (112 aa):
GSPELREKHRALAEQVYATWQELLKNTSNSPELREKLRALIEQVYATGQEMLKNGSVSPSPELREKHRALAEQVIATWQ
ELLKNTSNSPELREKFRALLEQVYATGQEMLKN
**DNA sequence:**

GGGAGTCCGGAACTACGTGAAAAACATCGGGCTTTGGCAGAGCAGGTTTATGCCACTTGGCAGGAACTTCTTAAG
AATACATCCAATAGCCCTGAGCTGCGTGAAAAATTACGTGCACTGATCGAACAAGTGTACGCCACGGGTCAAGAA
ATGTTGAAAAATGGCTCTGTAAGCCCCTCACCGGAACTGCGCGAAAAACACCGCGCCTTAGCGGAGCAGGTCATT
GCTACCTGGCAAGAACTGCTGAAAAACACCAGTAACTCGCCAGAACTCCGAGAGAAGTTTCGCGCGCTGCTGGAG
CAGGTGTATGCGACGGGCCAGGAAATGCTGAAAAAC

**>m4D2**
**Vector**: pET151
**AA sequence** (112 aa):
GSPELREKLRALIEQVYATGQEMLKNTSNSPELREKHRALAEQVYATWQELLKNGSVSPSPELREKFRALLEQVYATGQE
MLKNTSNSPELREKHRALAEQVIATWQELLKN
**DNA sequence:**
GGAAGTCCGGAACTTCGTGAAAAACTGCGTGCACTGATTGAACAGGTTTATGCAACCGGTCAGGAAATGCTGAAA
AATACGAGCAATAGCCCTGAGCTGCGCGAGAAACATCGCGCCCTGGCAGAGCAAGTCTACGCCACGTGGCAAGA
ACTGTTAAAGAACGGTAGCGTTTCTCCGTCACCAGAATTACGCGAAAAATTTCGGGCGCTTCTGGAACAAGTGTAT
GCCACAGGCCAAGAGATGCTTAAAAACACCTCGAACTCTCCTGAGCTGCGGGAAAAGCACCGTGCATTAGCCGAG
CAGGTTATTGCGACTTGGCAGGAATTACTGAAGAATTGA

**>4D2**
**Vector:** pET151
**AA sequence** (112 aa):
GSPELREKHRALAEQVYATGQEMLKNTSNSPELREKHRALAEQVYATGQEMLKNGSVSPSPELREKHRALAEQVYATG
QEMLKNTSNSPELREKHRALAEQVYATGQEMLKN
**DNA sequence**:
GGATCGCCAGAACTGCGCGAGAAACACCGTGCGTTAGCCGAACAAGTGTACGCCACAGGCCAAGAAATGCTGAA
GAACACGAGCAATTCGCCGGAACTTCGCGAGAAACATCGTGCTCTGGCAGAACAGGTGTATGCGACTGGCCAGG
AAATGCTGAAAAACGGGTCTGTAAGTCCGTCACCTGAACTGCGGGAGAAACACCGCGCTTTGGCCGAACAGGTTT
ACGCAACCGGTCAGGAGATGCTCAAGAACACCTCCAATAGCCCGGAACTGCGTGAGAAACATCGCGCATTAGCGG
AACAAGTCTATGCGACCGGTCAGGAAATGTTGAAAAAT

Where mentioned in the text, membrane protein designs were purified either with 10xHis or triple-StrepII tags at their C-termini, encoded by the DNA sequences below:

10xHis C-terminal tag, including a **triple alanine spacer**, **V5 epitope** and **10x His tag**

**AA sequence:**

**AAAGKPIPNPLLGLDSTHHHHHHHHHH**

**DNA sequence:**

GCGGCGGCTGGTAAACCGATCCCAAACCCTCTGCTTGGATTGGATTCCACACACCACCATCATCATCATCATCACCA
TCAT

Strep3 C-terminal tag, including a **triple alanine spacer**, **thrombin cleavage site** and **triple Strep-II tag**

**AA sequence:**

**AAA**LE**LVPRGSGGGSGGGSGGGSWSHPQFEKGGGSGGGSGGGSWSHPQFEKGGGSGGGSGGGSWSHPQFEK**

**DNA sequence:**

GCGGCCGCACTCGAGCTGGTTCCGCGTGGATCCGGTGGTGGGTCTGGTGGTGGGAGCGGTGGAGGCAGCTGGT
CGCATCCGCAGTTTGAGAAGGGCGGCGGATCAGGCGGCGGATCCGGCGGTGGCTCGTGGTCCCATCCGCAATTC
GAGAAGGGTGGCGGCAGTGGTGGCGGCTCTGGCGGTGGGTCGTGGAGCCACCCACAGTTCGAAAAG

**>CytbX**

**Vector:** pET-29

**AA sequence (113 aa):**

MGSPILRIIHLILALLVLITGLIMLLNTSNSPYLRLIHFLLALLVLITGWLMLKNGSKSPSPILRLIHIILAILVFITGIIMLLNTSNS
PFLRILHFILALLVFITGFLMLNQ

**DNA sequence:**

ATGGGCTCTCCTATTCTGCGCATCATTCACCTGATTTTGGCCTTGCTGGTTCTGATTACCGGACTTATCATGCTGCTG
AATACGTCAAATAGCCCCTATCTTCGCCTCATTCATTTTTTACTGGCACTGCTCGTGCTGATTACCGGTTGGCTGATG
CTAAAAAACGGTAGTAAGAGTCCGAGCCCGATCCTCCGTTTAATCCACATAATTCTGGCAATACTGGTATTTATTAC
TGGCATCATTATGTTACTGAACACATCGAACAGCCCATTCCTGCGGATTTTGCATTTCATCCTTGCGTTATTGGTCTT
TATCACGGGCTTCCTTATGCTGAACCAG

**>m1-CytbX (Rosetta decoy ID: 0016)**

**Vector:** pET-29

**AA sequence (113 aa):**

MGSPILRIIWLILLLLVLITGLIMLLNTSNSPYLRLIHFLLALLVLITFWLILKNGSKSPSPILRLIFIILIILVFITGIIMLLNTSNSPF
LRILHFILALLVMITAFLLLNQ

**DNA sequence:**

ATGGGCTCACCGATTCTGCGCATTATCTGGCTCATACTGCTTCTTCTGGTACTAATCACTGGCCTGATCATGCTTCTG
AATACCTCGAACAGCCCGTATCTGCGTCTGATTCACTTTCTGCTGGCATTATTAGTTCTGATTACCTTTTGGCTAATT
CTGAAAAATGGCTCTAAGAGCCCGTCGCCGATCCTGCGTTTGATTTTCATCATTCTCATTATTTTGGTCTTTATTACG
GGTATCATTATGCTGCTCAATACCAGTAACAGCCCCTTCCTGCGCATTCTGCACTTTATCCTCGCCTTGTTAGTGATG
ATTACGGCCTTCTTATTACTGAACCAG

**>m1-CytbX (Rosetta decoy ID: 0151)**
**Vector:** pET-29
**AA sequence (113 aa):**
MGSPILRIILLILFLLVLITGLIMLLNTSNSPYLRLIHFLLALLVLITLWLWLKNGSKSPSPILRLILIILLILVFITGIIMLLNTSNSPF
LRILHFILALLVLITFFLVLNQ
**DNA sequence:**
ATGGGCTCTCCCATTCTGCGCATAATTCTACTCATCTTGTTTCTTCTGGTATTAATTACCGGCCTGATTATGCTGTTA
AATACCTCAAACTCCCCGTATCTCCGTCTAATTCACTTTTTACTTGCACTGCTGGTGCTCATTACGCTGTGGCTGTGG
TTAAAAAATGGTAGCAAGTCGCCTTCGCCGATTCTGCGTTTGATCTTAATCATTCTGCTGATTTTGGTGTTCATCACT
GGCATCATTATGCTTCTGAACACCAGCAATAGTCCATTTTTGCGCATTCTCCACTTTATCCTGGCTCTTCTGGTCCTG
ATCACATTCTTCCTGGTTCTTAACCAG

**>m2-CytbX (Rosetta decoy ID: 0320)**
**Vector:** pET-29
**AA sequence** (113 aa):
MGSPILRIIHLILALLVLITFLIMLLNTSNSPYLRLILFLLMLLVLITGWLMLKNGSKSPSPILRLIHIILAILVFITIIILLLNTSNSPF
LRILLFILFLLVFITGFLMLNQ
**DNA sequence:**
ATGGGCTCCCCGATCCTGCGTATAATTCACCTCATCTTAGCGCTGTTGGTCCTGATTACATTTCTTATTATGTTGCTT
AACACGAGCAATTCTCCATATCTGCGTCTGATTCTTTTTCTGTTAATGCTGCTCGTACTGATTACCGGCTGGCTCATG
CTTAAGAATGGAAGCAAAAGCCCGAGTCCGATTCTGCGCCTAATTCATATTATCCTGGCGATTTTGGTGTTCATCAC
GATTATAATCCTGCTCTTGAATACCTCGAACTCGCCGTTTCTGCGCATCCTACTGTTCATTCTGTTCTTACTGGTTTTT
ATCACCGGTTTTCTGATGTTAAACCAG

**>m2-CytbX (Rosetta decoy ID: 0152)**
**Vector:** pET-29
**AA sequence (113 aa):**
MGSPILRIIHLILALLVLITFLIILLNTSNSPYLRLILFLLMLLVLITGWLMLKNGSKSPSPILRLIHIILAILVFITIIILLLNTSNSPFL
RILLFILFLLVFITGFLMLNQ
**DNA sequence:**
ATGGGTTCGCCTATCTTACGCATTATTCATCTTATCTTGGCACTGCTCGTGCTGATCACTTTTTTAATTATTCTGCTGA
ATACCTCCAATTCGCCGTATCTTCGTCTCATCTTATTCCTGCTGATGCTGTTGGTTTTGATCACCGGTTGGTTAATGC
TTAAAAACGGATCAAAAAGCCCGTCTCCGATTCTACGTCTGATACACATCATCCTAGCCATTCTGGTCTTTATTACC
ATTATTATTCTGCTGTTAAATACGAGCAACAGCCCGTTTTTACGCATTCTGCTGTTCATCCTCTTTCTTCTGGTATTTA
TTACAGGCTTCCTGATGCTCAACCAG

**>CytbX with m4D2 core packing mutations**
**AA sequence:**
MGSPILRIILLILILLVLITGLIMLLNTSNSPYLRLIHFLLALLVLITWWLLLKNGSKSPSPILRLIFIILLILVFITGIIMLLNTSNSPF
LRILHFILALLVIITWILLLNQ
**No DNA sequence was ordered**
